# Cell-specific FUS loss enhances ageing-associated transcriptional elongation in lung fibrosis

**DOI:** 10.64898/2026.10.07.757436

**Authors:** Diana G. Rogel-Ayala, Julio Cordero, Héctor A. Aguirre-Alarcón, Karla Rubio, Guruprasadh Swaminathan, Naceur Charif, Natalia de Isla, Witold Szymanski, Gowthami Kolluru Venkataswamy Reddy, Peter Braubach, Samina Mahmood, Stefan Günther, Miguel N. Rivera, Johannes Graumann, Thomas Braun, Malgorzata Wygrecka, Gergana Dobreva, Guillermo Barreto

**Affiliations:** Université de Lorraine, CNRS, Laboratoire IMoPA, UMR 7365, F-54000 Nancy, France; Lung Cancer Epigenetics, Max-Planck-Institute for Heart and Lung Research, 61231 Bad Nauheim, Germany; Department of Cardiovascular Genomics and Epigenomics, European Center for Angioscience (ECAS), Medical Faculty Mannheim, Heidelberg University, 68167 Mannheim, Germany; German Centre for Cardiovascular Research (DZHK), 68167 Mannheim, Germany; Massachusetts General Hospital and Harvard Medical School, Charlestown, MA 02129, USA; International Laboratory EPIGEN, Consejo de Ciencia y Tecnología del Estado de Puebla (CONCYTEP), Instituto de Ciencias, EcoCampus, Benemérita Universidad Autónoma de Puebla, 72570 Puebla, Mexico; Institute of Translational Proteomics & Core Facility Translational Proteomics, Department of Medicine, Philipps-Universität Marburg; 35043 Marburg, Germany; Center for Infection and Genomics of the Lung (CIGL), Faculty of Medicine, Justus Liebig University (JLU), Giessen, Germany; Institute for Lung Health (ILH), Justus Liebig University (JLU), Giessen, Germany; Member of the German Center for Lung Research.; Institute for Pathology, Hannover Medical School, Hanover, Germany; Biomedical Research in Endstage and Obstructive Lung Disease Hannover (BREATH) Research Network, Hanover, Germany; ECCPS Bioinformatics and Deep Sequencing Platform, Max-Planck-Institute for Heart and Lung Research, 61231 Bad Nauheim, Germany; Department of Cardiac Development, Max-Planck-Institute for Heart and Lung Research, 61231 Bad Nauheim, Germany; Center for Cancer Research, Massachusetts General Hospital and Harvard Medical School; Charlestown, MA, 02129, USA; Helmholtz-Institute for Translational Angio-CardioScience (HI-TAC) of the Max Delbrück Center for Molecular Medicine in the Helmholtz Association (MDC) at Heidelberg University, 69117 Heidelberg, Germany

**Keywords:** Fused in sarcoma (FUS), translocation speed of elongating RNA polymerase II, R-loops, cellular senescence, scRNA-seq, spatial transcriptomic, transcription regulation

## Abstract

Idiopathic pulmonary fibrosis (IPF) is an age-related, lethal lung disease affecting over 3 million people worldwide. During aging, impairment of diverse molecular processes compromises cellular homeostasis. Eukaryotic cells adapt to this ageing-associated stress by large-scale transcriptome changes affecting the expression of genes involved in signaling, protein homeostasis, cell plasticity and immune response. The molecular mechanisms regulating extensive transcriptome rearrangements leading to senescence in specific cell types of IPF lungs remain sparely investigated. Here we uncover fundamental molecular mechanisms underlying cellular senescence by integrative analysis of multi-omics studies and functional experiments monitoring fibrotic and senescence hallmarks in primary lung fibroblasts from control and IPF patients. Loss-of-function of the DNA/RNA binding protein fused in sarcoma (FUS) in control fibroblasts enhanced fibrotic and senescence hallmarks resembling IPF fibroblasts. Dissecting the molecular mechanisms, reduced FUS levels in IPF fibroblasts abolished R-loop formation at transcription start sites (TSS), leading to increased RNA polymerase II (Pol II) recruitment and its phosphorylation at serine 2, resulting in enhanced translocation speed of elongating Pol II and increased expression of fibrosis-and senescence-related genes. Summarizing, FUS maintains low Pol II processivity in control fibroblasts by stabilizing R-loops at TSS, acting as a gatekeeper of fibrotic and age-associated transcriptional programs.

## Main text

Transcription occurs in mechanistically distinct phases, including initiation, promoter-proximal pausing, elongation, and termination. Transition through these phases is regulated by ordered assembly of nucleic acid-protein complexes linking transcription to RNA processing. Phase-specific phosphorylation states of the carboxy terminal domain (CTD) of the largest Pol II subunit (RPB1) influence the interaction of the Pol II holoenzyme with different molecular complexes ^1^. Phosphorylation of serine 5 (S5p) is associated with initiation, promoter-proximal pausing, and recruitment of capping and early RNA-processing factors. In contrast, serine 2 phosphorylation (S2p) progressively increases along the gene body and is linked to productive elongation, co-transcriptional splicing, and 3′ end processing ^2,3^. During transcription initiation, the Pol II holoenzyme binds the promoter of genes to be transcribed and is released into elongation under various activating conditions, including S2p on the CTD of RPB1 ^1^. The transition from Pol II S5p to Pol II S2p depends, among others, on P-TEFb (Positive Transcription Elongation Factor b) activity and is a key checkpoint for releasing paused Pol II into elongation. FUS modulates this checkpoint by limiting premature S2p near the TSS ^4^, avoiding excessive or early release of Pol II into elongation. Interestingly, increased translocation speed of elongating Pol II together with alterations in pre-mRNA processing events have been related to aging in different species including humans ^5^.

IPF is the most common interstitial lung disease showing a incidence of 20 new cases per 100,000 persons per year ^6^. While the pathogenesis of IPF is complex involving multiple cell types, a central event is activation, proliferation and migration of fibroblasts following repetitive microinjuries of unknown origin to alveolar epithelial cells. This results in the formation of fibroblastic foci, which consist of highly proliferative fibroblasts, immune cells and excessive deposition of extracellular matrix (ECM) causing alterations of the alveolar framework, destruction of lung epithelium, stiffening of functional lung tissue and loss of gas exchange functionality. Current treatments alleviate patients’ symptoms and delay disease progression. Consequently, patients with IPF die within four years of diagnosis, primarily due to respiratory failure. Although the precise causes of IPF remain unknown, several risk factors have been identified, among which ageing plays a major role as the prevalence of IPF increases sharply with age. We have previously shown in primary human lung fibroblasts (hLF) isolated from IPF patients (IPF hLF) that epigenetic gene silencing and nucleolar organization mediated by the multicomponent RNA–protein complex MiCEE is compromised by hyperactive histone acetyltransferase EP300 (E1A binding protein p300), promoting transforming growth factor beta (TGF-β) signaling, fibrotic gene expression programs as well as fibroblast activation ^7,8^. Remarkably, we found FUS acting upstream of these mechanisms by inhibiting EP300 activation in hLF from control donors (Ctrl hLF). Thus motivated, this work delves into these findings by investigating the role of FUS in IPF.

### FUS loss in Ctrl hLF induces fibrotic hallmarks resembling IPF hLF

Total transcriptome analysis in Ctrl and IPF hLF revealed 6,652 differentially expressed genes (DEGs), from which 62.4% (n=4,148) were decreased and 37.4% (n=2,504) increased in IPF hLF as compared to Ctrl hLF (Fig. 1a; Extended Data Fig. 1a; Source Data file). Whereas established fibrotic markers, including *SNAI1*, *COL4A1*, *FN1*, *VIM*, *COL3A1* and, *COL1A1*, were among significantly increased transcripts in IPF hLF, *FUS* levels were significantly decreased (fold change = 0.7; *P*=0.001). This reduction was confirmed on the protein level (Fig. 1b-c). Interestingly, immunostaining showed FUS in the nucleus of Ctrl hLF excluding nucleoli. To demonstrate causal involvement of FUS in lung fibrosis we determined levels of fibrotic markers, such as FN1, COL1A1 and ACTA2 on RNA and protein levels (Fig. 1d-e; Extended Data Fig. 1a-c; Source Data file), and monitored collagen deposition and cell migration in Ctrl and IPF hLF (Fig. 1f-g; Extended Data Fig. 1d). We found that *FUS*-specific loss-of-function (LOF) by small interfering RNA (siRNA) in Ctrl hLF (further referred to as Ctrl hLF + FUS LOF) increased levels of fibrotic markers, collagen deposition and cell motility, as compared to Ctrl hLF transfected with Ctrl siRNA, mimicking the results obtained in IPF hLF. Further, FUS gain-of-function (GOF) after transfection of *FUS* expression construct into IPF hLF (further referred to as IPF hLF + FUS GOF) significantly reduced fibrotic markers and hallmarks as compared to IPF hLF and to similar levels as in Ctrl hLF. These results support a causal involvement of *FUS* LOF in lung fibrosis.

**Fig. 1:**
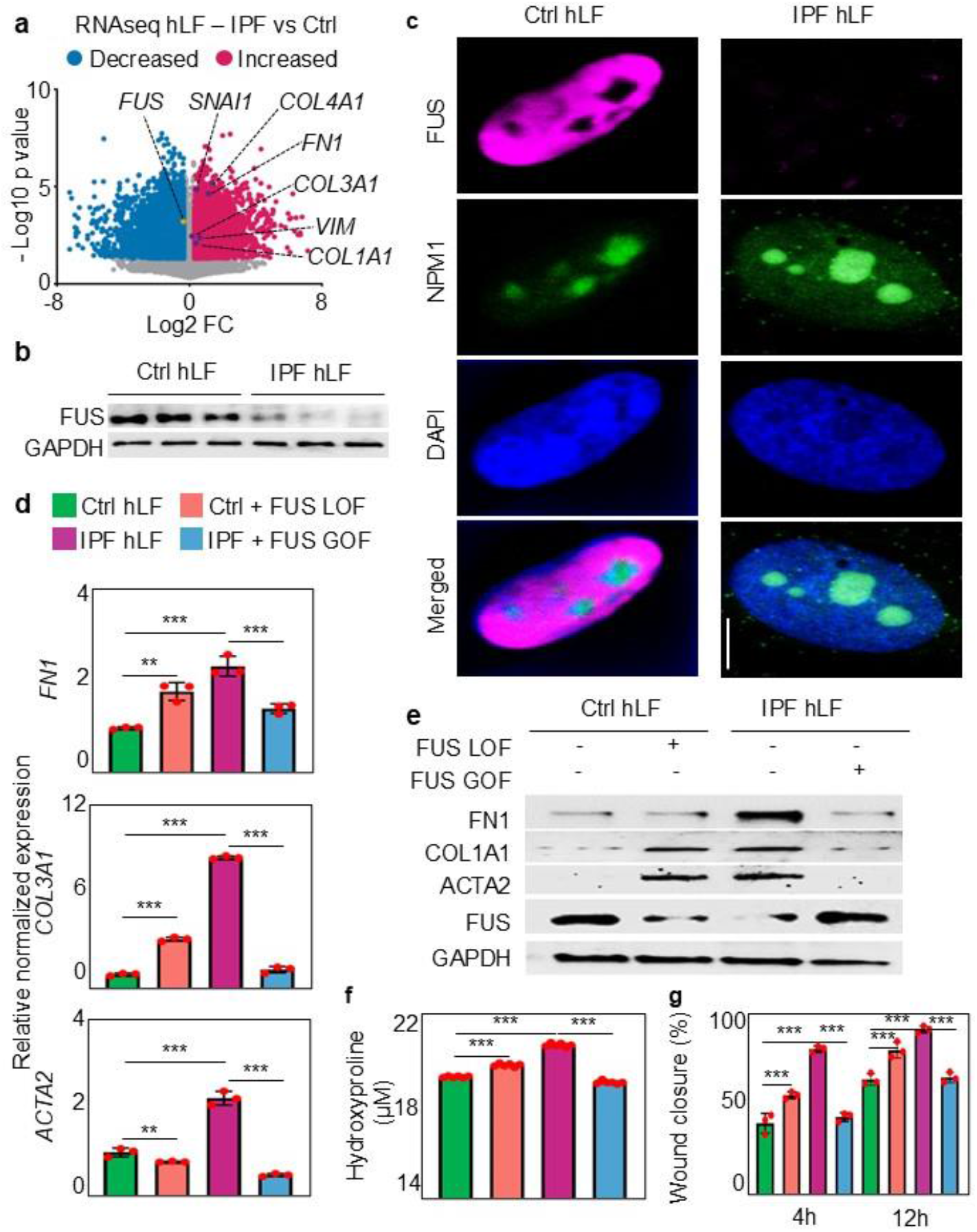
**FUS loss in Ctrl hLF induces fibrotic hallmarks resembling IPF hLF**. **a**, Volcano plot showing differentially expressed genes from RNA-seq of IPF hLF versus Ctrl hLF. Significantly decreased (blue) and increased (magenta) transcripts are shown. *FUS* is highlighted in yellow and fibrotic markers in lila. **b**, Western blot analysis of FUS in Ctrl hLF and IPF hLF (n = 3 per group). GAPDH was used as a loading control. **c**, Representative immunofluorescence images showing FUS (magenta) and the nucleolar marker NPM1 (green) in Ctrl hLF and IPF hLF. Nuclei were counterstained with DAPI (blue). Scale bar, 10 µm. **d**, RT-qPCR analysis of *FN1*, *COL3A1* and *ACTA2* expression in Ctrl hLF, Ctrl hLF + FUS LOF, IPF hLF and IPF hLF + FUS GOF (*n* = 3 biological replicates). **e**, Western blot analysis of FN1, COL1A1, ACTA2 and FUS in the indicated conditions. GAPDH was used as a loading control. **f**, Quantification of hydroxyproline levels in the indicated conditions (*n* = 6 biological replicates). **g**, Quantification of wound closure at 4 h and 12 h after scratch induction (*n* = 3 biological replicates). Bars represent mean ± SEM. *P* values by one-way ANOVA followed by Tukey’s post hoc test for multiple comparisons; \*\**P* < 0.01, \*\*\**P* < 0.001. See also Extended Data Fig. 1. Source data are provided as a Source Data file.

### Low FUS levels correlate with cell type-specific increased senescence

To place FUS in the pathological context of IPF, we analyzed publicly available spatial transcriptomic data in lung tissue of control and IPF patients ^9^, generated using the Visium technology for formalin-fixed and paraffin-embedded (FFPE) tissue sections for an almost whole-transcriptome wide mRNA analysis. From eight different cellular tissue microenvironments, or niches, with specific cellular composition found, three were related to IPF: fibrotic niche, airway macrophage niche, and immune niche ^9^. We focus our analysis on the fibrotic niche composed of myofibroblasts, aberrant basaloid cells, terminal bronchial secretory cells (TB-SCs), basal cells, and *IL1B+* macrophages (Fig. 2a-c; Extended Data Fig. 2-3; Source Data file). Most of the cells in the fibrotic niche showed low *FUS* expression (Fig. 2b; Extended Data Fig. 2-3), with a more pronounced effect in myofibroblast, aberrant basaloid cells, and *IL1B*+ macrophages (Fig. 2c). In addition, reduction of FUS in fibrotic lung tissue of IPF patients was confirmed on the protein level (Fig. 2d), supporting the results obtained by spatial transcriptomic analysis.

**Fig. 2:**
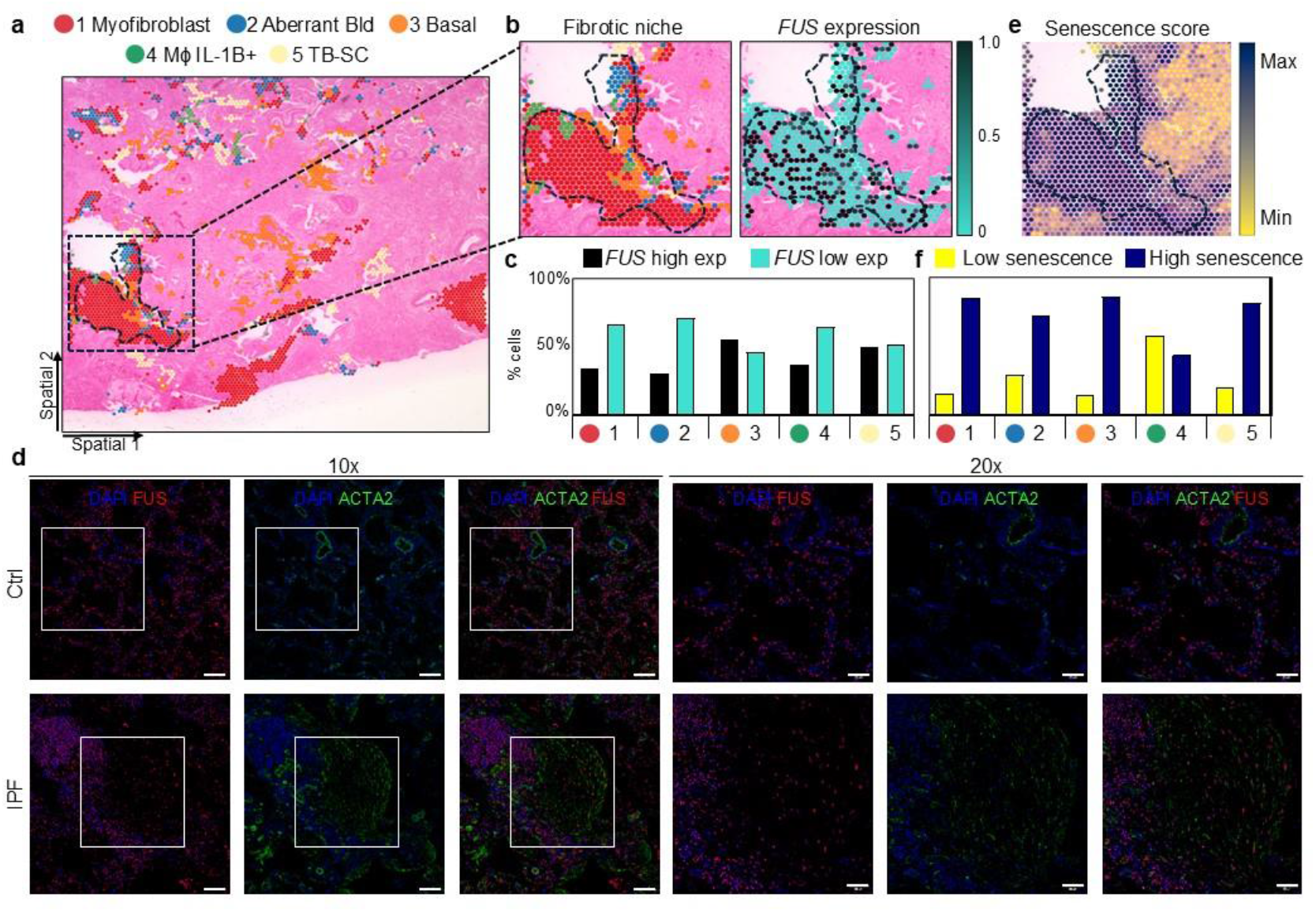
Low FUS levels correlate with increased cellular senescence. **a**, Spatial transcriptomic map of an IPF lung section showing five cell populations defining fibrotic niches ^9^: myofibroblasts, aberrant basaloid cells, basal cells, IL-1B+ macrophages and TB-SC cells. Dashed lines delineate a fibrotic niche. **b** and **e**, Magnified view of the indicated fibrotic niche showing cell populations, *FUS* expression and (**e**) senescence scores. **c** and **f**, Percentage of cells with high or low FUS expression and (**f**) low or high senescence scores within each cell population. **d**, Representative immunofluorescence microscopy pictures from three independent experiments showing FUS (red) and ACTA2 (green) in Ctrl and IPF lung tissue. DAPI, nuclei (blue). Squares are shown at higher magnification. Scale bars, 100 μm (left) and 50 μm (right). See also Extended Data Fig. 2-3. Source data are provided as a Source Data file.

IPF is an age-related disease characterized by hyperproliferation of specific fibroblast s in the fibrotic niche, even though myofibroblasts and aberrant basaloid cells (a recently identified, IPF-related cell state) have been associated with cellular senescence pathways ^10–12^. We therefore calculated a custom senescence gene score based on the expression of senescence markers in the spatial transcriptomic data and visualized it across the tissue sections ^13,14^ (Fig. 2e-f, Extended Data Fig. 3). Focusing on the fibrotic niche (Fig. 2e), the senescence gene score was high in the highlighted regions of cells with low *FUS* expression. Moreover, myofibroblasts, aberrant basaloid cells, basal cells and TB-SC with low *FUS* expression showed high senescence gene score, whereas the same cell types with high *FUS* expression displayed low senescence gene score (Fig. 2f). These results together supported a correlation between low *FUS* expression and high senescence gene score in cells of the fibrotic niche, which was confirmed using tissue sections from three additional IPF patients (Extended Data Fig. 3).

To complement results obtained by spatial transcriptomics, we harnessed the potential of single-cell RNA sequencing (scRNA-seq) to gain further cell-type-specific insight into pathological changes in IPF. To that end, three previously published scRNA-seq datasets from human lungs of Ctrl donors and IPF patients ^10,11,15^ were integrated to achieve greater sequencing depth per cell type. The integrated scRNA-seq datasets represent gene expression profiles of 224,532 cells from 50 human individuals (controls, n=30; IPF, n=20). Cell annotation was performed based on established single-cell signatures in the human lung ^11^, and represents 15 cell populations, characterized by distinct marker gene expression profiles (Fig. 3a; Extended Data Fig. 4; Source Data file). Focusing on the fibroblast population, we further annotated six fibroblast subtypes based on specific markers (Fig. 3b; Extended Data Fig. 5a-b), including resident alveolar fibroblasts, profibrotic fibroblasts, myofibroblasts, inflammatory fibroblasts, mesothelial-like fibroblasts and transition fibroblasts. We determined the proportion of each fibroblast subtype in control and IPF lungs (Fig. 3 c-d). Fibroblast subtypes related to lung fibrosis, such as profibrotic fibroblasts, myofibroblasts and mesothelial-like fibroblasts, were significantly over-represented in IPF patients as compared to Ctrl donors, whereas resident alveolar fibroblasts were significantly reduced (Fig. 3d). Interestingly, all the fibroblast subtypes showed significantly higher levels of the SenMayo gene expression signature (further referred to as SenMayo signature, a curated set of cellular senescence-associated genes validated across multiple tissues) ^13^, suggesting enhanced cellular senescence in IPF patients as compared to Ctrl donors (Fig. 3e). Remarkably, the IPF-related fibroblast subtypes, profibrotic fibroblasts, myofibroblasts and inflammatory fibroblasts, displayed significantly lower *FUS* expression in IPF patients as compared to Ctrl donors (Fig. 3f), whereas only resident alveolar fibroblasts from IPF patients showed higher *FUS* levels, which we will comment on the discussion. These results obtained from the integrated scRNA-seq datasets from human lungs of Ctrl donors and IPF patients ^10,11,15^ confirmed the correlation between low *FUS* expression and high senescence gene score in cells of the fibrotic niche previously determined by spatial transcriptomics.

**Fig. 3:**
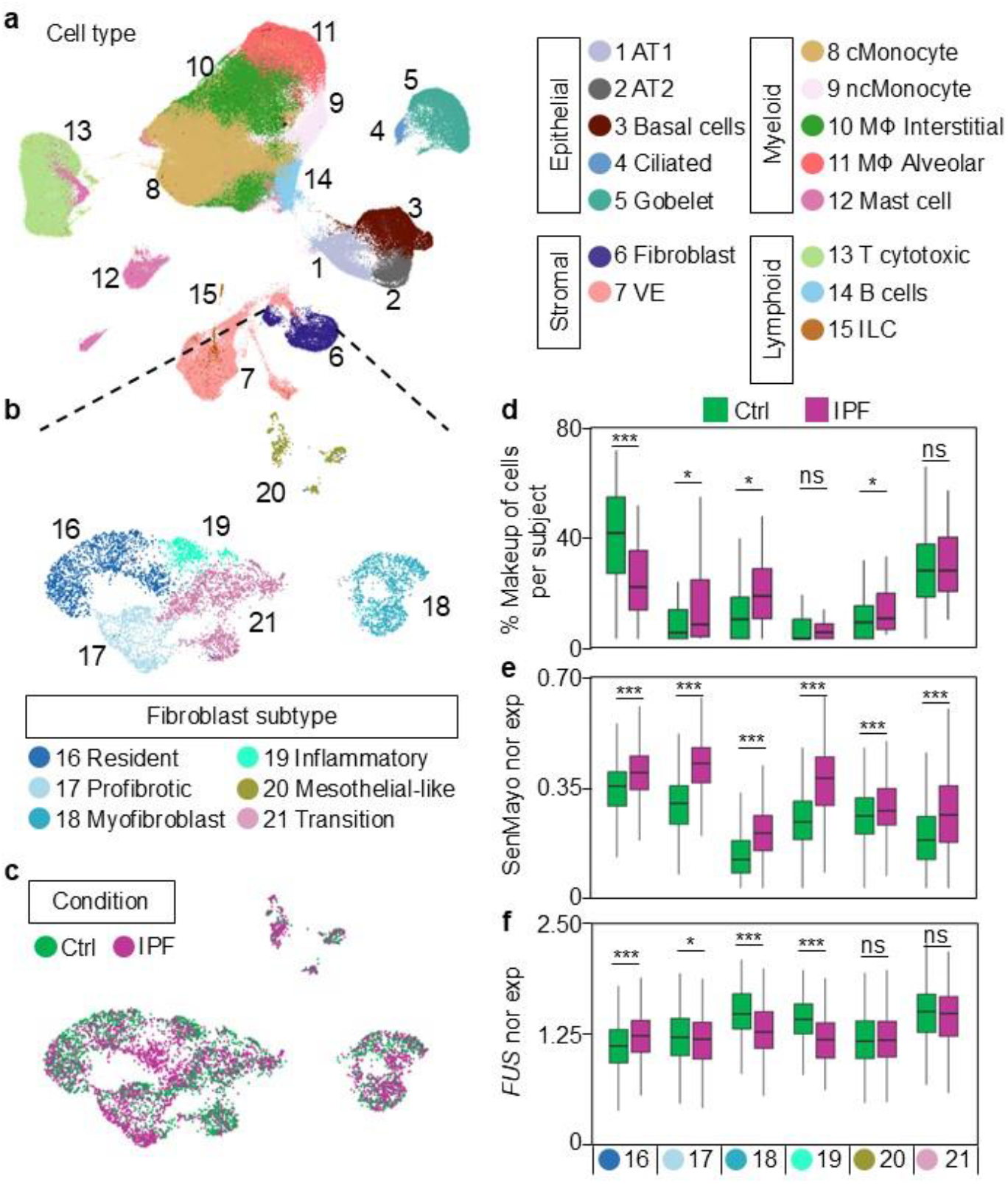
Low FUS expression is associated with increased SenMayo signature in IPF fibroblast subtypes. **a**, UMAP (Uniform Manifold Approximation and Projection) visualization of integrated scRNA-seq datasets showing the major epithelial, stromal, myeloid and lymphoid cell populations. **b**, Subclustering of fibroblasts into resident, profibrotic, myofibroblast, inflammatory, mesothelial-like and transition fibroblasts. **c**, UMAP coloured by condition, showing Ctrl (green) and IPF (magenta) fibroblasts. **d**, Boxplots showing the percentage of each fibroblast subtype per subject in Ctrl and IPF lungs. **e** and **f**, Boxplots showing normalized SenMayo gene signature expression (**e**) and FUS expression (**f**) across fibroblast subtypes in Ctrl and IPF lungs. Box plots indicate median (middle line), 25th, 75th percentile (box) and 5th and 95th percentile (whiskers). *P* values by two-tailed Wilcoxon rank-sum test; \**P* < 0.05, \*\*\**P* < 0.001, ns, non-significant. See also Extended Data Fig. 4-5. Source data are provided as a Source Data file.

### Cell-specific association of low FUS levels with increased SenMayo signature and Pol II speed in IPF

To further leverage the resolution obtained from the integrated scRNA-seq datasets ^10,11,15^, we first identified the IPF-specific cell niches ^9^: fibrotic niche, airway macrophage niche, and immune niche (Fig. 4a, Extended Data Fig. 6; Source Data file). As expected, the cell frequency in each of the IPF-specific cell niches was higher in IPF patients as compared to Ctrl donors, whereas in non-fibrotic cell niches the cell frequency was lower in IPF patients. Moreover, levels of the senescence-related SenMayo signature were increased in IPF patients in all the analyzed cell niches, suggesting pathology-related cellular senescence throughout all cellular niches (Fig. 4b). Interestingly, we detected significantly reduced *FUS* levels in the fibrotic, immune and non-fibrotic niches (Fig. 4c), supporting the observed correlation between reduced *FUS* levels and increased expression of senescence-related genes in these cellular niches, whereas cellular senescence in the airway macrophage niche is probably FUS-independent.

**Fig. 4:**
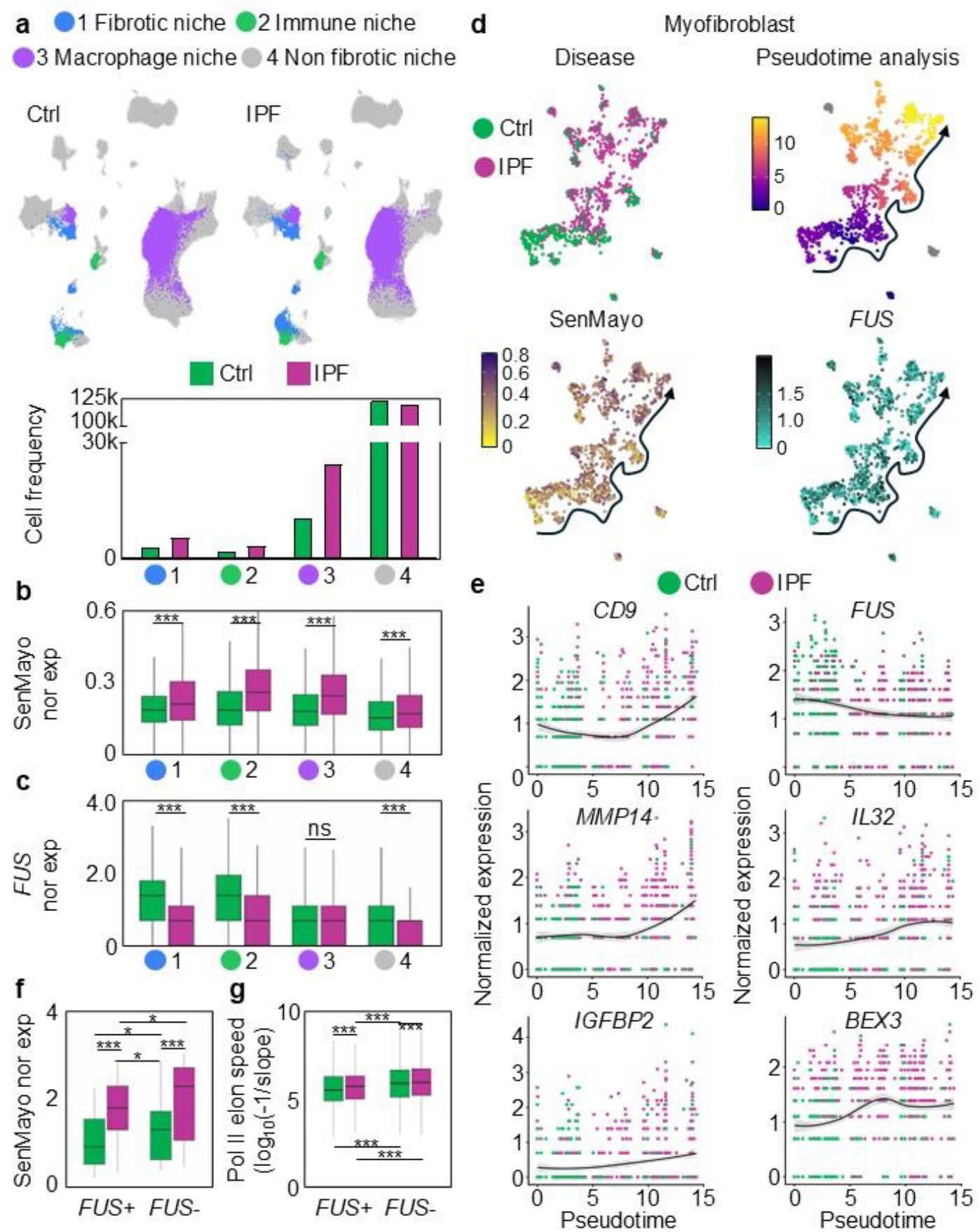
FUS expression decreases along IPF-associated myofibroblast pseudotime and correlates with increased SenMayo signature and enhanced Pol II elongation speed. a, UMAP visualizations of Ctrl and IPF lung cells classified into fibrotic, immune, macrophage and non-fibrotic niches ^9^, with the corresponding cell frequencies shown below. b and c, Boxplots showing normalized SenMayo gene signature expression (b) and *FUS* expression (c) across spatial niches in Ctrl and IPF lungs. d, UMAP visualizations of Ctrl and IPF myofibroblasts showing disease condition, pseudotime, SenMayo gene signature expression and *FUS* expression. e, Smoothed expression of selected genes along pseudotime trajectories in Ctrl and IPF myofibroblasts. f and g, Boxplots showing normalized SenMayo gene signature expression (f) and Pol II elongation speed (g) in fibroblasts with (*FUS*+) or without (*FUS*-) detectable *FUS* expression from Ctrl and IPF lungs. All box plots indicate median (middle line), 25th, 75th percentile (box) and 5th and 95th percentile (whiskers). *P* values by two-tailed paired Wilcoxon test (**b-c**) or two-tailed Student’s t-test (**f-g**); \**P* < 0.05, \*\*\**P* < 0.001, ns, non-significant. See also Extended Data Fig. 6-7. Source data are provided as a Source Data file.

To assess developmental progression and branching relationships of cells within myofibroblasts in the fibrotic niche, we determined diffusion pseudotime (DPT) distances ^16^ (Fig. 4d; Extended Data Fig. 7a-b). Unsupervised ordering of cells resulted in a clear separation of cells originating from Ctrl and IPF lungs (Fig. 4d; top left). Spearman correlation between DPT distances and gene expression revealed gradual increases in expression of genes commonly associated with activated myofibroblasts (Fig. 4d, top right; Extended Data Fig. 7a-b) with the highest expression of the respective genes at the IPF edge of each manifold. These results suggest a continuous trajectory toward IPF archetypes co-occurring within myofibroblasts. A similar gradual increase in expression of the SenMayo signature was observed with the highest expression at the IPF edge of each manifold (Fig. 4d, bottom left; Fig. 4e). Remarkably, the opposite pattern was observed for *FUS* expression, with the highest expression at the Ctrl edge and the lowest expression at the IPF edge of each manifold (Fig. 4d, bottom right; Fig. 4e). These results support the observed correlation between reduced *FUS* levels and increased expression of senescence-related genes in disease-related cell types of the IPF-specific fibrotic niche. To quantify these results, we separated cells of the integrated scRNA-seq datasets from human lungs of Ctrl donors and IPF patients ^10,11,15^ into cells with (*FUS*+) or without (*FUS*-) *FUS* expression and monitored the levels of the SenMayo signature (Fig. 4f). In both *FUS*+ and *FUS*− cells, the SenMayo signature was significantly higher in IPF than in control lungs, consistent with increased cellular senescence in IPF. Remarkably, in *FUS*-cells, levels of the SenMayo signature were significantly higher in both Ctrl and IPF cells as compared to the respective FUS-expressing Ctrl and IPF cells, demonstrating that the absence of *FUS* expression further enhanced cellular senescence-related gene expression.

Age-related increased Pol II elongation speed together with alterations in pre-mRNA processing events have been reported in various species including humans ^5^. Thus, as described in the Methods and commented in the Discussion, we adapted a method previously used to determine the translocation speed of elongating Pol II (further referred to as Pol II speed) in bulk RNA-seq using RNA-seq coverage in introns ^5,17,18^ to measure cell type-specific Pol II speed in the integrated scRNA-seq datasets from human lungs of Ctrl donors and IPF patients (Fig. 4g; Extended Data Fig. 7c). In both *FUS*+ and *FUS*-cells, we detected significantly higher Pol II speed in cells from IPF lungs as compared to Ctrl lungs, correlating with the IPF-related enhanced cellular senescence detected by the SenMayo signature. Moreover, in *FUS*-cells, Pol II speed was significantly increased in both Ctrl and IPF cells as compared to the respective *FUS*+ Ctrl and IPF cells, confirming the absence of *FUS* expression to further increase Pol II speed. Our results indicate that FUS keeps Pol II at a low-processivity rate acting as transcriptional break of gene expression programs associated to cellular senescence.

### FUS represses transcription of senescence-related genes

Due to the reduction of FUS in the nucleus of hLF of IPF patients (Fig. 1b-c), we decided to investigate this cell type more closely. We first assessed senescence-related transcriptional programs by determining the levels of specific gene sets using total RNA-seq (Fig. 5a; Extended Data Fig. 1a-c; Source Data file). Compared to Ctrl hLF, genes belonging to the senescence-related SenMayo signature showed significantly increased expression in both IPF hLF and Ctrl hLF + FUS LOF (Fig. 5a, top). Furthermore, senescent cells undergo extensive and context-dependent metabolic remodeling, which can include mitochondrial dysfunction, altered oxidative phosphorylation (OXPHOS) and increased production of reactive oxygen species (ROS) ^19,20^. Consistent with such metabolic remodeling, we detected significantly increased expression of transcripts belonging to the OXPHOS gene set in both IPF hLF and Ctrl hLF + FUS LOF compared with Ctrl hLF (Fig. 5a, middle). In addition, in IPF hLF and Ctrl hLF + FUS LOF we found significant reduction of genes regulating the cell cycle (Fig. 5a, bottom), in line with another defining feature of senescent cells, their stable cell cycle arrest. Remarkably, overexpression of FUS in IPF hLF was insufficient to revert the senescence-related transcriptional programs detected in IPF hLF. However, to complement the results obtained by transcriptomic analysis, we monitored hallmarks of cellular senescence in Ctrl hLF and IPF hLF (Fig. 5b-d; Extended Data Fig. 8; Source Data file). CDKN1A (further referred to as p21) is a cyclin-dependent kinase inhibitor inducing cell cycle arrest in senescent cells ^21^ and one of the commonly used senescent cell markers. As compared to Ctrl hLF, we detected significantly increased number of p21-positive (p21+) hLF in IPF and in Ctrl hLF + FUS LOF (Fig. 5b; Extended Data Fig. 8a). Interestingly, FUS GOF in IPF hLF significantly reduced the number of p21+ cells as compared to IPF hLF. Another stablished cellular senescence marker is the cytochemical detection of endogenous, pH 6-active, senescence-associated beta-galactosidase (SA-β-gal) ^22^, an enzyme that is active in most cells at pH 4, indicating their normal state. Correlating with p21 results, we detected increased SA-β-gal in IPF hLF and in Ctrl hLF + FUS LOF as compared to Ctrl hLF (Fig. 5c; Extended Data Fig. 8b). In contrast to p21 results, FUS GOF in IPF hLF did not significantly change SA-β-gal as compared to IPF hLF. Loss of cellular proliferation is another characteristic of cellular senescence and can be monitored by labeling newly synthesized DNA using 5-Ethynyl-2‘-deoxyuridine (EdU). As compared to Ctrl hLF, whereas we detected significantly reduced cell proliferation in IPF hLF, no significant differences in cell proliferation were observed in Ctrl hLF + FUS LOF (Fig. 5d;

**Fig. 5:**
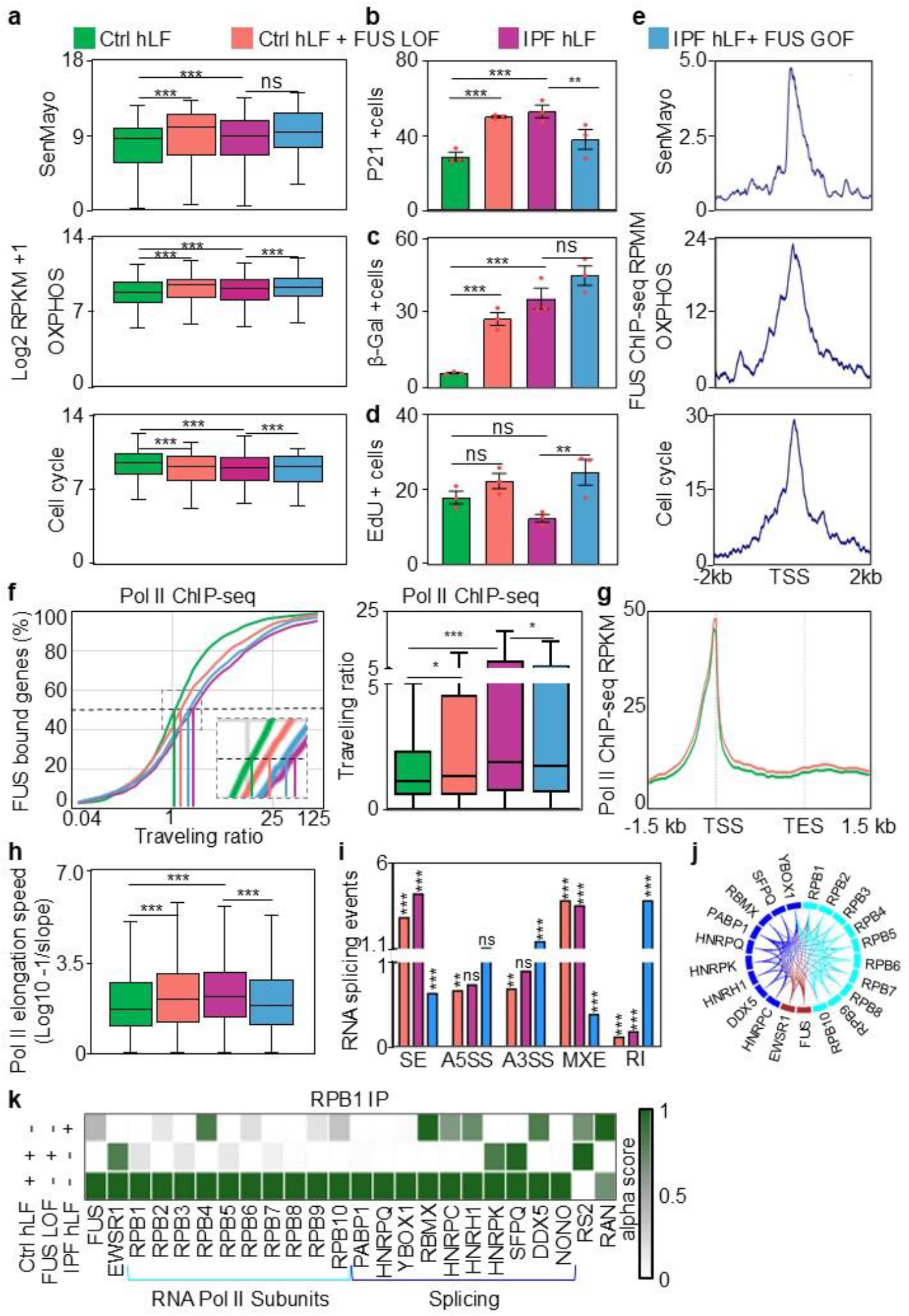
FUS represses transcription of senescence-related genes. **a**, Boxplots showing expression of genes belonging to the SenMayo, OXPHOS and cell-cycle signatures in Ctrl hLF, Ctrl hLF + FUS LOF, IPF hLF and IPF hLF + FUS GOF. Reads were normalized using RPKM (reads per kilobase of transcript per million mapped reads) and represented as log2. **b**-**d**, Quantification of P21-positive (**b**), β-galactosidase-positive (**c**) and EdU-positive (**d**) hLF as in (**a**) (*n* = 3 biological replicates). **e**, Aggregate plots showing FUS enrichment by ChIP-seq at the transcription start sites (TSS) ± 2kb of genes belonging to the SenMayo, OXPHOS and cell-cycle signatures. **f**, Cumulative distribution of the Pol II travelling ratio for FUS-bound genes and corresponding boxplots of travelling-ratio determined in Pol II ChIP-seq in hLF as in (**a**). **g**, Pol II ChIP-seq profiles from the TSS to the transcription end site (TES). **h**, Boxplots showing Pol II elongation speed (log10[1/slope]) in hLF as in (**a**). **i**, Ratios of major alternative splicing event frequencies relative to the corresponding reference condition: skipped exon (SE), alternative 5′ splice site (A5SS), alternative 3′ splice site (A3SS), mutually exclusive exon (MXE) and retained intron (RI). **j**, STRING protein-protein interaction network centred on RPB1, showing Pol II subunits, FUS and splicing regulators. **k**, RPB1 immunoprecipitation followed by mass spectrometry in Ctrl hLF, Ctrl hLF + FUS LOF and IPF hLF. Alpha scores indicate the relative abundance of co-immunoprecipitated proteins. Bars represent mean ± SEM. All box plots indicate median (middle line), 25th, 75th percentile (box) and 5th and 95th percentile (whiskers). *P* values by two-tailed paired Wilcoxon test (**a**), one-way ANOVA followed by Tukey’s post hoc test (**b**-**d**, **f** and **h**) or two-tailed Student’s t-test (**i**); \**P* < 0.05, \*\**P* < 0.01, \*\*\**P* < 0.001, ns, non-significant. See also Extended Data Fig. 8. Source data are provided as a Source Data file.

Extended Data Fig. 8b). Our results monitoring senescence-related transcriptional programs and hallmarks of cellular senescence demonstrated increased cellular senescence in hLF when FUS levels were reduced, namely in IPF hLF and Ctrl hLF + FUS LOF. To further link FUS to these findings, we analyzed a publicly available sequencing experiment subsequent to chromatin immunoprecipitation (ChIP-seq) using FUS-specific antibodies ^4^. We detected FUS enrichment at transcription start sites (TSS)-/+ 2 kb of the genes belonging to the senescence-related gene sets that we assessed by total RNA-seq (SenMayo, OXPHOS and cell cycle gene sets; Fig. 5e), suggesting a role of FUS in transcriptional regulation of senescence-related genes.

FUS plays an important role in transcription by binding the CTD of RPB1 and inhibiting S2p at TSS of genes bound by FUS (further referred to as FUS targets) ^4,23^, thereby controlling the release of Pol II into the elongation phase. Thus, we decided to determine the “Pol II traveling ratio” ^24^ using total Pol II ChIP-seq for calculating the ratio of the density of Pol II near the TSS (-300/+100 nucleotides to TSS) over that in the gene body of FUS targets (Fig. 5f; Source Data file). We observed a significant increase in the median Pol II traveling ratio of FUS targets in IPF hLF (2.0; *P*=7.05e-5) and Ctrl hLF + FUS LOF (1.3; *P*=0.038) as compared to Ctrl hLF (1.1). Furthermore, we detected significantly higher accumulation of Pol II at the TSS and in gene bodies of FUS targets in Ctrl hLF + FUS LOF as compared to Ctrl hLF (Fig. 5g), supporting an increase in Pol II recruitment to FUS targets, which together with the increased expression of the same FUS targets (Fig. 5a) suggest an enhanced Pol II activity at these genes. We further measured Pol II speed using RNA-seq coverage in introns ^5,17^. Read coverage of total RNA-seq typically declines 5′ to 3′ along an intron. The magnitude of this decrease reflects Pol II speed: the milder the reduction, the faster the elongation ^5,18^. We detected significantly increased Pol II speed in IPF hLF and Ctrl hLF + FUS LOF as compared to Ctrl hLF (Fig. 5h; Extended Data Fig. 9a; Source Data file), correlating with increased Pol II travelling ratios (Fig. 5f) and enhanced cellular senescence (Fig. 5a-d) in hLF with reduced *FUS* levels (IPF hLF and Ctrl hLF + FUS LOF), as well as with the age-related increased Pol II speed reported in different species ^5^. Remarkably, FUS GOF in IPF hLF significantly reduced the median Pol II travelling ratios of FUS targets (1.69; *P*=0.013) and Pol II speed (1.79; *P*=1.26e-12) as compared to IPF hLF (1.83 and 2.16, respectively), supporting a partial rescue of these processes since it is not reflected in the expression of senescence-related genes (Fig. 5a).

Transcription elongation is crucial for proper mRNA synthesis, as pre-mRNA processing steps, including RNA splicing, occur in parallel to transcription ^25^. Indeed, malfunction of transcriptional elongation results in the formation of erroneous transcripts and may lead to diseases ^26^. Thus, we quantified the relative abundance of five primary categories of RNA splicing events (skipped exons, SE; alternative 5′ splice sites, A5SS; alternative 3′ splice sites, A3SS; mutually exclusive exons, MXE; and retained introns, RI) in our RNA-seq in hLF (Fig. 5i; Extended Data Fig. 9b). Compared to Ctrl hLF, hLF with reduced *FUS* levels (IPF hLF and Ctrl hLF + FUS LOF) displayed significant increase in SE and MXE events, whereas A5SS, A3SS and RI events were reduced. Interestingly, FUS GOF in IPF hLF counteracted the effects observed in IPF hLF, leading to significant reductions in SE and MXE events, whereas A3SS and RI events were significantly increased. To further investigate the role of FUS in transcription regulation and RNA splicing, we first determined the protein-protein interaction network of RBP1 using the biological data base STRING ^27^ (Fig. 5j) and confirmed the interaction with FUS and further Pol II subunits. In addition, we found EWSR1, another member of the FET protein family ^28^, as well as various proteins involved in RNA splicing interacting with RBP1. Furthermore, we performed a high-resolution mass spectrometry-based proteomic analysis following immunoprecipitation of RBP1 from protein extracts of Ctrl hLF and confirmed these protein-protein interactions (Fig.5k; Source Data file). RS2 and RAN were used as negative controls. Moreover, most of these interactions were weakened in IPF hLF and Ctrl hLF + FUS LOF suggesting that FUS is required for the adequate formation of the Pol II holoenzyme and protein complexes mediating RNA splicing.

### FUS stabilizes R-loops at TSS and represses fibrotic-and senescence-related genes

RPB1 precipitated proteins related to R-loops in FUS-dependent manner (Fig. 5k) ^29^. We thus decided to investigate whether these three-stranded nucleic acid structures consisting of a DNA-RNA hybrid and a displaced single strand of DNA ^30^ play a role in transcriptional regulation of fibrotic-and senescence-related genes mediated by FUS (Fig. 6). Since FUS binds both DNA and RNA ^28^, we compared FUS-specific ChIP-seq ^4^ and CLIP-seq (Cross-Linking Immunoprecipitation followed by Sequencing) ^31^ and found 7,150 genes/transcripts common to both data sets (Fig. 6a; Source Data file). In addition, we determined the genome-wide distribution of R-loops in Ctrl hLF by CUT&Tag-seq (Cleavage Under Targets and Tagmentation followed by Sequencing; Extended Data Fig. 10a). Focusing on the TSS of the 7,150 genes common between FUS-specific ChIP-seq and CLIP-seq, unsupervised clustering of the CUT&Tag signal defined four distinct clusters based on R-loop enrichment (Fig. 6b; Source Data file). Genes in cluster 1 (C1; n = 1,313) and cluster 2 (C2; n = 1,190) displayed R-loop enrichment 3‘ (upstream) of the TSS spanning the windows –2 kb to –0.5 kb and – 0.5 kb to TSS, respectively. Genes in cluster 3 (C3; n = 1,135) and cluster 4 (C4; n = 2,602) showed R-loop enrichment 5‘ (downstream) of the TSS within the windows TSS to +0.5 kb and +0.5 kb to +2 kb, respectively. We also performed R-loop-specific CUT&Tag in Ctrl hLF + FUS LOF and IPF hLF and found that independent of the position of the R-loop relative to the TSS, FUS is required for the formation of the R-loop (Fig. 6c). Visualization of the loci of selected FUS targets using the integrative genomic viewer (IGV) confirmed these results in fibrotic and senescence-related genes (Fig. 6d; Extended Data Fig. 10b). To gain further insight into these findings, we performed a high-resolution mass spectrometry-based proteomic analysis following immunoprecipitation of DNA–RNA hybrids using the S9.6 antibody (further referred to as R-loops IP) from protein extracts of Ctrl and IPF hLF (Fig. 6e; Source Data file). R-loops IP in Ctrl hLF precipitated a broad repertoire of proteins spanning several functional categories. These included classical R-loop-related proteins (DDX21, DDX5, DHX9, NONO, and SFPQ), alternative splicing factors (HNRNPC, HNRNPH1, HNRNPL, and RBMX), splicing regulators (ROA2, HNRPK, HNRPM, HNRPR, HNRPU, NUCL, and NOP2) and ribosomal RNA processing proteins (RL11, RL13, RL14, and RL5). AKAP8 and PHAX were presented as negative controls. The presence of these diverse functional categories of proteins underscores the functional integration of R-loops into different aspects of RNA processing in Ctrl hLF. Remarkably, the precipitation of these proteins by R-loops IP was weakened in Ctrl hLF + FUS LOF and IPF hLF supporting the requirement of FUS for the interaction of R-loops with these proteins.

**Fig. 6:**
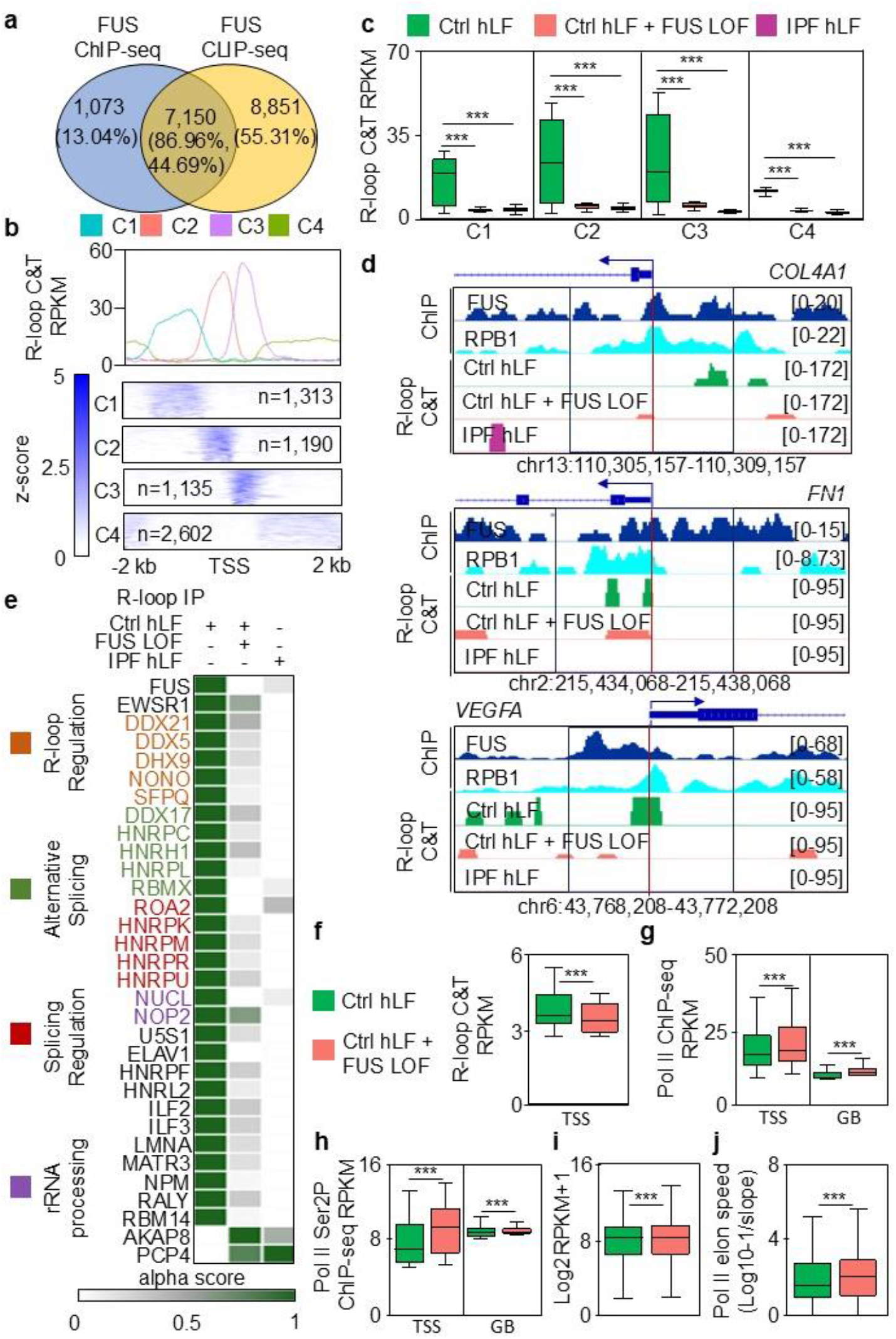
FUS negatively regulates fibrotic-and senescence-related genes by stabilizing R-loops at TSS. **a**, Venn diagram showing the overlap between genes/transcripts bound by FUS ChIP-seq and FUS CLIP-seq. **b**, Enrichment plots after R-loop-specific CUT&Tag at transcription start sites (TSS) ± 2 kb of FUS targets, grouped by unsupervised clustering into four R-loop enrichment patterns: C1 (−2 kb to −500 bp), C2 (−500 bp to the TSS), C3 (TSS to +500 bp) and C4 (+500 bp to +2 kb). Average enrichment profiles (top) and heatmaps (bottom) are shown for each cluster. **c**, Boxplots showing R-loop enrichment by CUT&Tag across clusters C1-C4 in Ctrl hLF, Ctrl hLF + FUS LOF and IPF hLF. **d**, Genome browser tracks showing FUS ChIP-seq, RPB1 ChIP-seq and R-loop CUT&Tag signals at the loci of fibrosis-related genes. R-loop tracks are shown for Ctrl hLF, Ctrl hLF + FUS LOF and IPF hLF. Red vertical lines indicate TSS, and signal ranges are shown on the right. **e**, Immunoprecipitation of DNA-RNA hybrids using the S9.6 antibody followed by proteomic analysis in Ctrl hLF, Ctrl hLF + FUS LOF and IPF hLF. Detected proteins are grouped as R-loop regulators, alternative splicing factors, splicing regulators and rRNA-processing proteins. Alpha scores indicate the relative abundance of co-immunoprecipitated proteins. **f-j**, Boxplots showing enrichment at TSS or gene bodies (GB) of FUS targets of R-loops by CUT&Tag (**f**), total Pol II (**g**) and Pol II S2p (**h**) by ChIP-seq, gene expression of FUS targets by RNA-seq (**i**), and Pol II elongation speed (log10[1/slope]) at FUS targets in Ctrl hLF and Ctrl hLF + FUS LOF. All box plots indicate median (middle line), 25th, 75th percentile (box) and 5th and 95th percentile (whiskers). In **a**-**c** and **d**-**i**, values were normalized using reads per kilobase per million (RPKM). *P* values by one-way ANOVA followed by Tukey’s post hoc test (**c**) or two-tailed Student’s t-test (**f-j**); \*\*\**P* < 0.001, ns, non-significant. See also Extended Data Fig. 9-10. Source data are provided as a Source Data file.

To correlate the results on R-loops with the results on Pol II, we determined the enrichment of R-loops, total Pol II and transcription elongating Pol II S2p at the TSS of FUS targets with canonical Pol II S2p enrichment pattern across the gene ^4^ in Ctrl hLF and Ctrl hLF + FUS LOF (Fig. 6f-h). Whereas R-loop enrichment at TSS was significantly reduced upon FUS LOF as compared to Ctrl cells (Fig. 6f), levels of total Pol II (Fig. 6g, left) and Pol II S2p (Fig. 6h, left) significantly increased, suggesting enhanced Pol II recruitment to the TSS and release from promoter-proximal pausing into transcriptional elongation once the TSS have been cleared from R-loops. We also detected increased total Pol II (Fig. 6g, right) and Pol II S2p (Fig. 6h, right) enrichment in the gene bodies of FUS targets following FUS LOF, which together with the increased expression of the same FUS targets (Fig. 6i) support enhanced Pol II activity at these genes. In line with these interpretations, we measured significantly higher Pol II speed at the same FUS targets in Ctrl hLF + FUS LOF as compared to Ctrl hLF (Fig. 6j).

## Discussion

Our results at the cellular level, which support FUS as inhibitor of pathological fibroblast activation and cellular senescence, are explained at the molecular level through further results demonstrating a mechanism, in which FUS is required for R-loop formation at TSS of fibrotic and senescence-related genes acting as transcriptional break of gene expression programs associated to fibrosis and cellular senescence. In cells with low FUS levels, such as IPF hLF or Ctrl hLF + FUS LOF, the promoters of these genes are cleared of FUS and R-loops allowing an enhanced recruitment of total Pol II and increased S2p, which in turn releases Pol II from promoter proximal-pausing into transcriptional elongation. In addition, (1) increased Pol II traveling ratio, (2) augmented Pol II speed and (3) enhanced expression of fibrotic and senescence-related genes, together support enhanced Pol II activity at these genes. On the other hand, in hLF from control donors, FUS acts as a gatekeeper of fibrotic and age-associated transcriptional programs by maintaining low Pol II processivity at these genes, stabilizing R-loops and inhibiting Pol II S2p at TSS. The connection between FUS and Pol II kinetics is supported by *in vitro* reconstitution assays showing that FUS binds peptides containing the CTD of RPB1 and blocks S2p, as well as ChIP-seq in HEK293T cells showing increased enrichment of Pol II S2p upon FUS LOF at TSS of genes with canonical Pol II S2p pattern ^4,32,33^. Even considering differences between experimental systems used in these previous studies, the convergence of the observations with our results support a conserved role for FUS in tuning Pol II elongation kinetics. Importantly, we placed these findings in the pathological context of an age-related disease (IPF) leveraging the potential of cutting-edge technologies, such as spatial transcriptomics and scRNA-seq, to demonstrate in a FUS-dependent manner,cell type-specific augmented Pol II speed correlating with increased levels of the senescence-associated SenMayo signature.

A recent study reported increased total FUS expression and cytoplasmic accumulation in primary interstitial fibroblasts from patients with IPF ^34^. This apparent discrepancy with our results may reflect fibroblast heterogeneity, as our single-cell RNA-seq analyses show that reduced FUS expression and increased cellular senescence occur in specific fibroblast subtypes rather than across the entire fibroblast population. Consequently, bulk analyses of isolated fibroblasts may mask these cell type-specific differences by averaging FUS expression across heterogeneous populations.

Interestingly, whereas FUS binding to CTD was RNA-dependent, the S2p inhibition was not ^4^, suggesting a role of R-loops as proposed here. R-loops are intimately connected to early co-transcriptional RNA processing, influence splice-site selection, and serve as recruitment hubs for RNA helicases, RNA-binding proteins, and splicing regulators. In this context, FUS is ideally positioned to support R-loop stability: it binds RNA with high affinity, interacts with nascent transcripts at promoter-proximal regions, and associates with regulatory complexes (including NONO/SFPQ, hnRNPs and DDX-family helicases) that modulate R-loop formation and spliceosome assembly ^35,36^. Our R-loops IP in Ctrl hLF recovered a coherent set of RNA-binding factors and R-loop-associated proteins with established roles in resolving or stabilizing R-loops, promoting splice-site recognition, or linking nascent RNA to higher-order ribonucleoprotein assemblies. The fact that these R-loop-protein interactions were reduced in cells with low FUS levels (IPF hLF and Ctrl hLF + FUS LOF) indicates that FUS is required for assembling or maintaining these R-loop-protein complexes shaping the promoter-proximal molecular environment.

Genome-wide analyses in multiple species have shown average Pol II elongation speed to increase with age ^5^. Motivated by these findings, we extended the use of scRNA-seq beyond its conventional applications by determining Pol II elongation speed. We applied this framework to three previously published scRNA-seq datasets from human lungs of Ctrl donors and IPF patients ^10,11,15^. Although 3′ scRNA-seq protocols are designed to capture polyadenylated transcripts, multiple studies have reproducibly shown approximately 20% of the reads to align to intronic regions ^37–40^, reflecting pre-mRNA presence and incomplete processing rather than technical noise. We used the intronic reads of the integrated scRNA-seq datasets from human lungs of Ctrl donors and IPF patients ^10,11,15^ to quantify Pol II elongation speed based on intronic read coverage, a strategy previously applied only in bulk or single-nucleus RNA-seq ^5,17,18^. Standard workflows, such as those implemented in Seurat ^41^, primarily rely on gene-level quantification of spliced, polyadenylated transcripts, and discard intronic reads by default. This practice is rooted in the assumption that intronic reads carry little biological meaning in single-cell datasets, particularly in 10x Genomics v2/v3 protocols that emphasize polyA enrichment. Consequently, analyses that depend on intronic information (such as transcriptional kinetics, alternative splicing, or co-transcriptional processing) have historically been avoided in scRNA-seq. The idea that Pol II speed may be inferred from scRNA-seq has been considered technically difficult due to the expected scarcity of introns, an obstacle we overcame in this study by integrating three different scRNA-seq datasets ^10,11,15^ to achieve greater sequencing depth per cell type, improve reproducibility, and increased statistical power. The consistency in our results of intronic signal across conditions and fibroblast subtypes further supports its biological relevance rather than being a technical artifact. Thus, although unconventional, our analyses are methodologically justified by properties inherent to the datasets, as well as by growing evidence that intronic reads in scRNA-seq contain interpretable information about nascent transcription and co-transcriptional processing. Finally, the integrative framework presented here may be strengthened in future work using orthogonal approaches. Both single-nucleus RNA-seq and scRNA-seq using Parse Biosciences (using Evercode combinatorial barcoding) contain a significantly higher proportion of intronic reads as compared to standard 10x Genomics scRNA-seq libraries, offering ideal options to validate our results demonstrating cell type-specific enhanced Pol II speed correlating with increased SenMayo signature in cells with low FUS levels. In addition, nuclear run-on assays (such as GRO-seq, PRO-seq, or TT-seq) may be another option to confirm the results obtained here using 10x Genomics scRNA-seq, since they directly measure nascent transcription rates by tracking active RNA polymerases. Together, our approach addresses a conceptual gap in the scRNA-seq field using the biological information contained in the 20% of the intronic reads inherent of the datasets generated with scRNA-seq technologies implementing polyA enrichment (such as in 10x Genomics v2/v3 protocols), thereby opening a door to use the vast and growing universe of data generated with scRNA-seq technologies to perform cell type-specific analyses of transcriptional kinetics, alternative splicing, or co-transcriptional processing.

## Methods

### Cell culture

Primary human lung fibroblasts (hLF) were obtained from patients with idiopathic pulmonary fibrosis (IPF hLF) or from non-fibrotic lung tissue resected from patients undergoing surgery for lung cancer (Ctrl hLF). Fibroblasts were isolated at the Center for Infection and Genomics of the Lung, Justus Liebig University Giessen, using the outgrowth method. All patient-derived material was anonymized. Cells were cultured in MCDB-131 medium (Gibco, #10372019) supplemented with 8% fetal calf serum, 1% L-glutamine, 1% penicillin–streptomycin (100 U/mL each), 2 ng/mL fibroblast growth factor, 0.5 ng/mL epidermal growth factor, and 5 µg/mL insulin, as previously described ^8^. Cells were maintained at 37 °C in a humidified incubator containing 5% CO₂ and were routinely tested for mycoplasma contamination. For subculturing, cells were washed with 1× PBS and detached using 0.25% trypsin–EDTA. All experiments were performed using cells below passage 10. MRC5 was grown in EMEM (ATCC, 30-2003) All media were supplemented with 10% FBS and cells were cultured at 37 °C with 5% CO2. Cells were maintained and split every 2–4 days according to ATCC recommendations.

### Cell transfection and FUS loss-and gain-of-function

Cells were transfected using Lipofectamine 2000 (Invitrogen) following the manufacturer’s instructions. For FUS loss-of-function, Ctrl hLF were transfected with MISSION® esiRNA targeting human FUS (Sigma-Aldrich/Merck, EHU092111) at a final concentration of 50 nM. For FUS gain-of-function, IPF hLF were transfected with 4 µg of FLAG-tagged full-length human FUS expression plasmid (FLAG-fl-hFUS) together with 0.5 µg of empty vector per 10-cm culture dish. A total of 9 µL of Lipofectamine 2000 was used per dish.

### Total RNA isolation and RNA sequencing data analysis

Total RNA was isolated from human lung fibroblasts using TRIzol reagent (Invitrogen) according to the manufacturer’s instructions. Briefly, cells were lysed directly in TRIzol, followed by phase separation with chloroform. The aqueous phase was collected, and RNA was precipitated with isopropanol at −20 °C overnight. After centrifugation, RNA pellets were washed with 75% ethanol, resuspended in RNase-free water, and quantified using a NanoDrop spectrophotometer (Thermo Fisher Scientific). Total RNA sequencing was performed using two biological replicates for each experimental condition. Libraries were sequenced in paired-end mode on a NextSeq 500 instrument (Illumina) using V2 chemistry.

Raw sequencing reads were assessed using FastQC v0.11.8. Adapter sequences and low-quality reads were removed using Trimmomatic v0.27 with the following parameters: ILLUMINACLIP:${ADAPTERS}:2:30:10 LEADING:3 TRAILING:3 SLIDINGWINDOW:4:15 HEADCROP:5 MINLEN:15. High-quality reads were aligned to the human reference genome GRCh38/hg38 using STAR v2.7.9 and GENCODE v39 gene annotations. SAM files were converted to BAM format using Samtools v1.12. Tag directories were generated using makeTagDirectory from HOMER v4.11.1, and gene expression was quantified as reads per kilobase per million mapped reads using analyzeRepeats.pl with the command analyzeRepeats.pl rna hg38-count genes-rpkm.

To avoid division by zero during fold-change calculations, genes with an RPKM value of zero were assigned a minimum value of 0.0001. Differential expression was assessed for IPF hLF versus Ctrl hLF, Ctrl hLF with FUS loss-of-function versus Ctrl hLF, and IPF hLF with FUS gain-of-function versus IPF hLF. Genes with a log₂ fold change ≥ 0.5 and *P* < 0.05 were considered upregulated, whereas genes with a log₂ fold change ≤ −0.5 and *P* < 0.05 were considered downregulated.

### Western blot analysis

Protein extracts were mixed with loading buffer, denatured at 95 °C, and separated by SDS– PAGE using 10% polyacrylamide gels. Proteins were transferred onto nitrocellulose membranes at 100 V for 1 h under chilled conditions. Successful transfer was verified by Ponceau S staining. Membranes were blocked with 5% milk in PBST and incubated overnight at 4 °C with primary antibodies against FUS (Santa Cruz Biotechnology, #sc-47711), fibronectin 1 (FN1; BD Biosciences, #610077), collagen type I alpha 1 (COL1A1; Rockland Immunochemicals, #600-401-103), α-smooth muscle actin (α-SMA; GeneTex/Biozol, #GTX124505-50), or GAPDH (Cell Signaling Technology, #97166), each diluted 1:1,000 in blocking solution. Where applicable, membranes were blocked with 5% bovine serum albumin for phosphoprotein detection.

The following day, membranes were incubated with the appropriate horseradish peroxidase-conjugated secondary antibodies diluted 1:10,000 and washed three times with PBST. Chemiluminescent signals were detected using WesternBright Sirius substrate (Advansta) and acquired with a ChemiDoc XRS imaging system (Bio-Rad).

### Immunofluorescence staining

Human lung fibroblasts were seeded onto ethanol-sterilized coverslips placed in 24-well plates and cultured until the desired cell density was reached. Cells were fixed with 4% paraformaldehyde for 15 min, washed with PBS, and permeabilized with 0.4% Triton X-100 in PBS. Non-specific binding was blocked using 3% bovine serum albumin and 0.1% Triton X-100 in PBS. Non-specific binding was blocked using 3% bovine serum albumin (BSA) and 0.1% Triton X-100 in PBS. Cells were incubated with primary antibodies against FUS (Santa Cruz Biotechnology, #sc-47711; 1:500), nucleophosmin 1 (NPM1; Santa Cruz Biotechnology, #sc-56622; 1:200), or α-smooth muscle actin (α-SMA; GeneTex/Biozol, #GTX124505-50; 1:200). After washing, cells were incubated for 1 h at room temperature in the dark with the appropriate fluorescent secondary antibodies: goat anti-rabbit IgG Alexa Fluor 488 (Thermo Fisher Scientific, #A-11008) or goat anti-mouse IgG Alexa Fluor 594 (Thermo Fisher Scientific, #A-11005), both diluted 1:1,000. Cells were subsequently washed three times with washing buffer and once with PBS. Coverslips were mounted onto glass slides using a DAPI-containing mounting medium and stored protected from light at 4 °C until imaging. Fluorescence images were acquired using a confocal microscope.

For p21 staining, cells grown on coverslips were fixed with 4% paraformaldehyde for 15 min, permeabilized with 0.5% Triton X-100 in PBS, and blocked using 0.5% BSA and 0.1% Triton X-100 in PBS. Cells were incubated with a primary antibody against p21 (Becton Dickinson, #556431; 1:250), followed by the appropriate fluorescent secondary antibody. Nuclei were counterstained with DAPI (Sigma-Aldrich), and fluorescence images were acquired using the Axio Scope.A1 microscope from Zeiss. Positive staining for p21 was quantified as the percentage of p21-positive cells relative to the total number of DAPI-positive cells. Three independent biological replicates were analyzed, with 10 fields quantified per replicate.

Human lung samples were fixed with 4% buffered formaldehyde, dehydrated, and paraffin-embedded. Sections (2.5 µm) were deparaffinized and subjected to heat-induced antigen retrieval in TRIS-EDTA buffer (10 mM Tris-HCl, 1mM EDTA, pH 9.0), followed by blocking with 3% bovine serum albumin (BSA) in TBS-T (20 mM Tris Base, 150 mM NaCl, 0.1% Triton X-100, pH 7,4) for 20 min at room temperature. Primary antibodies, mouse monoclonal anti-FUS (4H11; 1:50; Santa Cruz Biotechnology, Santa Cruz, CA, cat. no.: sc-47711) and mouse monoclonal anti-α-SMA labeled with Cy3 (1:300, Sigma-Aldrich, Darmstadt, Germany; cat. no.: C6198) diluted in 0.5% BSA in TBST were applied and the slides were sequentially incubated overnight at 4 °C. Corresponding, Alexa Fluor-conjugated secondary antibody was diluted 1:500 in 3% BSA in TBS-T and administered for 1 h at room temperature. Between the incubation steps, slides were washed with TBS-T buffer. Nuclear counterstaining was performed using 1 mg/ml DAPI solution (Thermo Scientific). Images were obtained with the Leica TCS-SP8 confocal microscope (Leica).

### Quantitative reverse transcription PCR

Quantitative PCR was performed using SYBR Green chemistry according to the manufacturer’s instructions. Reactions were carried out using a StepOne Real-Time PCR System (Applied Biosystems) and were performed in technical triplicates.

Data were analyzed using StepOne Software v2.3. Relative gene expression was calculated using the 2^−ΔΔCt method, with GAPDH as the reference gene. Primer sequences are provided in Extended Data Table 1.

### Hydroxyproline assay

Hydroxyproline levels were measured using the QuickZyme Hydroxyproline Assay Kit (QuickZyme Biosciences, #QZBhypro1) according to the manufacturer’s instructions. Briefly, cell extracts corresponding to 3 × 10⁶ cells per condition were mixed 1:1 (v/v) with 12 M HCl and hydrolyzed at 95 °C for 20 h. After cooling, samples were centrifuged at 13,000 × *g* for 10 min, and the supernatants were collected and diluted with distilled water.

Absorbance was measured at 570 nm using a microplate reader, and hydroxyproline concentrations were calculated from a standard curve generated using trans-4-hydroxy-L-proline provided with the kit. Results are expressed as µg/mL hydroxyproline. The experiment was performed using two biological replicates, each analyzed in three technical replicates.

### Wound closure assay

The wound closure assay was performed using cells cultured in 24-well plates. Once the cells reached confluence, a uniform scratch was created in the cell monolayer using a sterile pipette tip. Images of the wound area were acquired immediately after scratching (0 h) and at 4 and 12 h post-wounding.

Images were analyzed using ImageJ software, and the percentage of wound closure was calculated relative to the initial wound area at 0 h. The experiment was performed using three biological replicates.

### Spatial transcriptomics Analysis

Spatial transcriptomics data from idiopathic pulmonary fibrosis (IPF) lungs generated by Mayr et al. 2024 ^9^ were used in this study. The publicly available Visium dataset, including the processed gene expression matrix, spatial coordinates, tissue annotations, and previously defined spatial niches, was used as provided by the authors. Analyses focusing on the fibrotic niche were performed using the original niche annotations reported in the published dataset, without modification or reclassification. Analysis of the spatial Visium RNA-seq data was performed with the Python packages SCANPY (version 1.9.3) ^42^, Squidpy (1.2.3) ^43,44^, and cell2location (0.1.3) ^45^. Analysis of the spatial Visium dataset was performed with the Python packages SCANPY (version 1.9.3) ^42^, Squidpy (1.2.3) ^43,44^, and cell2location (0.1.3) ^45^.

To investigate the spatial distribution of *FUS* expression within selected cell populations, spots were identified based on cell-type-specific abundance thresholds. Thresholds were individually adjusted for each tissue section to account for section-specific differences in cell-type abundance distributions. Five cell populations were considered: myofibroblasts, aberrant basaloid cells, TB-SC cells, IL1B-positive macrophages, and basal cells. For each tissue section, spots exceeding the corresponding threshold for at least one of these cell populations were retained for downstream visualization. Based on *FUS* normalized expression, spots with values ≤0.6 were classified as *FUS* low expression, whereas spots with values >0.6 were classified as *FUS*-high expression. *FUS* expression was subsequently mapped onto the selected spatial spots using the spatial plotting framework implemented in SCANPY. Based on the average normalized expression of the senescence genes *SERPINE1, CDKN2B, IGFBP2, IGFBP7, SEMA3F, MMP14,* and *CD9*, spots with values ≤1.2 were classified as low senescence, whereas spots with values >1.2 were classified as high senescence.

### Senescence gene score

A senescence gene score was generated from the expression of *SERPINE1*, *CDKN2B*, *IGFBP2, IGFBP7, SEMA3F, MMP14,* and *CD9* using the compute_scores function from hotspot ^14^. Gene expression was modeled with a depth-adjusted negative binomial model, followed by gene-wise centering and smoothing across neighboring cells. Principal component analysis was then used to integrate the signature genes into a single score. The analysis was performed on the log1p counts layer using 30 neighbors, with the X_scVI embedding defining the neighborhood relationships ^9^. The resulting score was used to assess the spatial distribution of senescence.

### Sample preparation and Liquid Chromatography – Tandem Mass Spectrometry (LC-MS/MS)

Proteins bound to the magnetic beads were recovered through two successive elutions using 2% SDS in 50 mM Tris-HCl (pH 8.0) with incubation at 95 ° C for 10 min shaking and polled. Reduction of disulfide bridges was performed with 10 mM dithiothreitol (DTT; 95 ° C, 10 min), followed by carbamidomethylation of free sulfhydryl groups using 13 mM iodoacetamide (IAA) for 30 min at ambient temperature in the dark.

Subsequent clean-up and enzymatic proteolysis followed a modified single-pot solid-phase– enhanced sample preparation (SP3) protocol ^46^ utilizing a custom magnetic rack. Comprehensive parameters covering protein binding, on-bead digestion, and peptide recovery are deposited at https://doi.org/10.5281/zenodo.17474920. Resulting peptides were recovered in 45 µL of 0.1% formic acid containing 0.015% (w/v) n-dodecyl-β-D-maltoside (DDM) and transferred into autosampler vials. Peptide yields were determined fluorometrically (Pierce Fluorometric Peptide Assay, Thermo Scientific) to equalize mass load across runs.

Approximately 25 ng of peptide digest per sample was loaded onto a nanoElute liquid chromatograph system coupled with a timsTOF Ultra mass spectrometer (Bruker Daltonics). A PepMap C18 µ-Precolumn (5 mm, Thermo Scientific) was used, followed by separation across an analytical PepSep Ultra C18 column (75 µm × 25 cm, 1.5 µm particle size; Bruker Daltonics) at a continuous flow rate of 300 nL/min. Solvents consisted of 0.1% aqueous formic acid (A) and 0.1% formic acid in acetonitrile (B). Elution was achieved with a multi-step gradient: 5% to 10% B over 5 min, followed by an increase to 35% B over 25 min. The eluate was introduced into the mass spectrometer via a 20 µm CaptiveSpray emitter.

Mass spectra were acquired in dia-PASEF mode according to standard instrument configurations. Full-scan MS1 data were collected across an m/z range of 100–1,700 at a resolution of 45,000. Ion mobility separation was conducted over an inverse reduced mobility 1/K₀ window of 0.64–1.45 V·s/cm² using a 100 ms accumulation/ramp cycle. For DIA fragmentation, the precursor window spanned m/z 400–1,000 divided into 24 distinct isolation steps of 25 m/z each.

Detailed description of the LC as well as MS acquisition methods can be found here: https://doi.org/10.5281/zenodo.17475274.

### Proteomic data processing, visualization and protein interaction analysis

DIA files were processed in library-free workflow using DIA-NN (v2.0 Academia) ^47^ with deep learning-based in silico spectral generation against the canonical human UniProtKB/Swiss-Prot database (reviewed, November 2024, 20,429entries).

The digestion enzyme was set to trypsin (cleaving C-terminal to Lys/Arg, blocked by adjacent Pro) with a maximum of one missed cleavage. Search parameters included peptide lengths of 7–30 amino acids, precursor charge states of 1+ to 4+, and precursor/fragment m/z limits of 100–1,800. Carbamidomethylation of cysteine was configured as a static modification, whereas methionine oxidation (up to two sites per peptide) and N-terminal methionine loss were set as variable modifications. The precursor false discovery rate (FDR) was set at 1%, and the match-between-runs (MBR) algorithm was enabled.

Downstream statistical analysis was performed in R using the Autonomics package (v1.15.325) https://www.bioconductor.org/packages/release/bioc/html/autonomics.html. Protein identifications were filtered to maintain an FDR q-value threshold at 0.01. Protein groups lacking quantified precursors (n < 1) within a given replicate were filtered out, and remaining missing-not-at-random (MNAR) data were replaced via imputation in Autonomics by drawing values from a down-shifted, narrowed normal distribution. Raw, unnormalized intensities were log2-transformed and subsequently used for differential expression analysis performed using empirical Bayes moderated t-statistics via the limma pipeline ^48^.

Protein–protein interaction networks were generated using the STRING database based on experimentally determined and predicted interactions among selected proteins identified by mass spectrometry. Circos plots were generated in R using the circlize package (version 0.4.18) to visualize the relationships between selected proteins and their associated biological processes. Protein interactions from RPB1 IP were visualized using heatmaps generated with Morpheus (Broad Institute) based on normalized protein abundance values from the mass spectrometry dataset.

### CUT&Tag R-loop signal profiling at FUS-associated regions

Genomic regions associated with FUS binding were defined based on previously identified FUS targets from ChIP-seq and CLIP-seq datasets. R-loop CUT&Tag signal was evaluated at these FUS-associated regions using RPKM-normalized BigWig files. For each gene-associated region, a TSS ± 2 kb window was generated. The mean R-loop signal was calculated for each bin by integrating BigWig signal values across overlapping genomic intervals, generating a signal matrix representing the distribution of R-loop enrichment around FUS-associated TSS regions. Regions were grouped according to their R-loop signal profiles using k-means clustering (k = 4). Heatmaps were generated using the EnrichedHeatmap R package (version 1.40.1) to visualize R-loop enrichment patterns across clustered regions. Average signal profiles were calculated from the binned signal matrix and visualized in R.

### Single-cell RNA sequencing data analysis

Publicly available single-cell RNA-sequencing datasets of lung tissue from control donors and patients with idiopathic pulmonary fibrosis were obtained from the Gene Expression Omnibus under accession numbers GSE136831, GSE135893, and GSE214085. Samples were selected to obtain comparable control and IPF groups based on age, sex, ethnicity, and smoking status, resulting in 50 human individuals (30 controls and 20 IPF patients). Raw sequencing data were processed using Cell Ranger to generate gene–barcode expression matrices.

Gene–cell expression matrices were imported into R and analyzed using Seurat v5.4. The complete dataset comprised 224,532 cells (122,292 Ctrl and 102,240 IPF cells) before fibroblast sub-setting. Cells expressing fewer than 200 or more than 9,000 genes were excluded before normalization and downstream analysis. Expression data were normalized using SCTransform, followed by principal component analysis, sample integration, nearest-neighbour graph construction, unsupervised clustering at a resolution of 0.5, and UMAP visualization. Clusters were annotated according to the expression of canonical lineage markers.

Major lung cell populations were identified using established marker genes. Epithelial populations were annotated using markers including EPCAM, SFTPC, SCGB1A1, and FOXJ1; endothelial cells using PECAM1, VWF, and CLDN5; and immune cells using PTPRC together with lineage-specific markers. Fibroblasts were identified based on the expression of stromal markers, including PDGFRA, COL1A1, POSTN, and MFAP4, whereas myofibroblast identity was supported by the expression of ACTA2, TAGLN, and MYL9.

Fibroblast cells were subsequently subset from the integrated Seurat object and independently reprocessed by normalization, dimensionality reduction, nearest-neighbour graph construction, and unsupervised sub-clustering at a resolution of 0.3. The fibroblast subset comprised 7,991 cells, including 2,687 cells from control samples and 5,304 cells from IPF samples. Fibroblast clusters were classified into six transcriptionally distinct subtypes: resident, profibrotic, myofibroblast, transition, inflammatory, and mesothelial-like fibroblasts, based on the expression of subtype-specific marker genes shown in Extended Data Fig. 6a.

Normalized gene-expression values were used to evaluate FUS expression and gene-set expression across fibroblast subtypes. For each cell, senescence and fibrotic marker scores were calculated as the mean normalized expression of all genes included in the corresponding marker set, whereas FUS was analyzed using its individual normalized expression value. Cells were grouped according to fibroblast subtype and disease condition, and the resulting distributions were visualized using boxplots.

### Senescence-associated β-galactosidase staining and EdU incorporation assay

Senescence-associated β-galactosidase activity and cell proliferation were assessed 24 h after transfection using the Senescence β-Galactosidase Staining Kit (Cell Signaling Technology) and the Click-iT Plus EdU Alexa Fluor 594 Imaging Kit (Thermo Fisher Scientific), respectively, according to the manufacturers’ instructions. Senescence-associated β-galactosidase activity was detected at pH 6, with positive cells identified by the presence of blue cytoplasmic staining.

For proliferation analysis, cells were incubated with EdU for 48 h to label newly synthesized DNA. Total nuclei were counterstained with DAPI. Images were acquired using a Leica DMI3000B fluorescence microscope, with at least 15 representative fields analyzed per condition. β-galactosidase and EdU positivity were quantified as the percentage of positive cells relative to the total number of DAPI-positive cells. Cells positive for β-galactosidase staining and negative for EdU incorporation were classified as senescent. Experiments were performed using three independent biological replicates.

### Chromatin immunoprecipitation sequencing

Chromatin immunoprecipitation was performed using 4 × 10⁶ cells per immunoprecipitation. Cells were crosslinked with 1% formaldehyde for 10 min at room temperature, and the reaction was quenched with glycine at a concentration of 125mM. Cells were washed and collected in ice-cold PBS supplemented with protease and phosphatase inhibitors. Nuclei were isolated using hypotonic lysis buffer and resuspended in nuclear lysis buffer.

Chromatin was fragmented using a Covaris M220 sonicator, and insoluble material was removed by centrifugation. Sheared chromatin was pre-cleared with blocked beads for 2 h at 4 °C. For each immunoprecipitation, 25 µg of chromatin DNA was incubated with 2 µg of FUS antibody (Santa Cruz, #sc-47711) or control IgG (Santa Cruz, #sc-2027). overnight at 4 °C. Immunocomplexes were captured using blocked Protein G Sepharose beads and sequentially washed with low-salt, high-salt, LiCl, and TE buffers supplemented with protease and phosphatase inhibitors.

Immunoprecipitated DNA was eluted using an SDS-containing buffer. RNA and proteins were removed by treatment with RNase A and Proteinase K, respectively, and crosslinks were reversed overnight at 65 °C under high-salt conditions. DNA was purified using a commercial PCR purification kit and quantified using a NanoDrop spectrophotometer.

For MRC5 cell line Chromatin from 5M formaldehyde-fixed cells was fragmented to a size range of 200–700 bases with a Branson 250 Sonifier. Solubilized chromatin was immunoprecipitated with 5 μg antibodies against H3K27ac (Active Motif, 39133), Pol II Ser2Ph (Abcam, ab5095) and Pol II (Abcam, ab300575) at 4 °C overnight. Antibody– chromatin complexes were pulled down with protein G Dynabeads (Life Technologies, 10004D), washed, and then eluted. After cross-link reversal and RNase (Roche, 43813100) and proteinase K (Invitrogen, 25530049) treatment, immunoprecipitated DNA was extracted with AMP Pure beads (Beckman Coulter, A63881). ChIP DNA was quantified with Qubit dsDNA HS Assay kit (Invitrogen, Q32854). ChIP DNA samples were used to prepare sequencing libraries with Ultralow V2 DNA-Seq Library Preparation Kit (NuGEN, 0344NB-A01) and DNA samples were sequenced with the Nextseq 2000 Illumina genome analyzer.

### ChIP-seq data processing and peak annotation

Raw sequencing reads were assessed for quality, and high-quality reads were aligned to the human reference genome GRCh38/hg38 using Bowtie2 v2.4.2 with default parameters. SAM files were sorted and converted to BAM format using Samtools v1.12. PCR duplicates were removed using the MarkDuplicates function from Picard Tools v3.3.0.

Deduplicated BAM files were converted to BED format using Bedtools. Tag directories were generated using makeTagDirectory from HOMER v4.11.1 with default settings. RPKM-normalized coverage tracks were generated from the deduplicated BAM files using bamCoverage from deepTools v3.5.6.

Peak calling was performed using MACS2 v2.2.9.1 with the parameters callpeak-g hs-q 0.05 --broad --extsize 200 --keep-dup 1 --nomodel. The corresponding input samples were included as controls. Peaks overlapping ENCODE blacklisted regions were removed using bedtools intersect-v. The remaining peaks were assigned to genomic features and nearby genes using annotatePeaks.pl from HOMER.

### ChIP-seq enrichment analysis

ChIP-seq enrichment was evaluated using RPKM-normalized coverage tracks. Average occupancy profiles were generated to examine the distribution of FUS, total RNA polymerase II (RPB1), and RNA polymerase II phosphorylated at serine 2 across annotated gene regions from the transcription start site to the transcription end site.

FUS occupancy was additionally assessed around the transcription start sites of genes belonging to the SenMayo, oxidative phosphorylation, and cell-cycle gene sets. Gene coordinates were obtained from the hg38 annotation, and normalized ChIP-seq signal was summarized across promoter-centred regions to generate average enrichment profiles. These profiles were used to compare FUS enrichment at genes associated with senescence, mitochondrial metabolism, and cell-cycle regulation.

### FUS CLIP-seq analysis

Publicly available FUS CLIP-seq data from Jutzi et al. ^31^ were retrieved from the Gene Expression Omnibus under accession number GSE139263. Sequencing data were processed using a workflow adapted from nf-core/clipseq (https://github.com/nf-core/clipseq). Raw sequencing reads were assessed using FastQC v0.11.8. Adapter sequences and low-quality reads were removed using Trimmomatic v0.27 with the following parameters: ILLUMINACLIP:${ADAPTERS}:2:30:10 LEADING:3 TRAILING:3 SLIDINGWINDOW:4:15 HEADCROP:5 MINLEN:15. High-quality reads were aligned to the human reference genome GRCh38/hg38 using STAR. FUS-binding regions were identified by peak calling using MACS2. The resulting CLIP-seq peaks were assigned to genomic features and nearby genes using annotatePeaks.pl from HOMER. Genes associated with FUS CLIP-seq peaks were classified as FUS CLIP-seq-associated genes.

### FUS targets definition by integration of ChIP-seq and CLIP-seq analysis

FUS target genes were defined by integrating the FUS ChIP-seq and CLIP-seq datasets. FUS ChIP-seq-associated genes and FUS CLIP-seq-associated genes were identified independently as described above. Genes detected in both datasets were classified as FUS target genes. The intersection yielded 7,150 genes, which were used for downstream analyses of R-loop enrichment and RNA polymerase II transcriptional dynamics.

### RNA polymerase II travelling ratio analysis

The RNA polymerase II travelling ratio was calculated to assess the relative distribution of polymerase between promoter-proximal regions and gene bodies ^4^. For each annotated gene, the promoter-proximal region was defined as the interval extending from 30 bp upstream to 300 bp downstream of the transcription start site. The gene body was defined as the region extending from 300 bp downstream of the transcription start site to the transcription end site. RNA polymerase II read density was calculated from the ChIP-seq alignment files using bamCoverage from deepTools v3.5.6 and normalized as reads per kilobase per million mapped reads. The travelling ratio was calculated for each gene as the normalized RNA polymerase II read density within the promoter-proximal region divided by the normalized read density across the gene body. Genes shorter than 1 kb or showing insufficient ChIP-seq coverage were excluded from the analysis.

Travelling ratios were calculated for FUS-bound genes and compared among Ctrl hLF, Ctrl hLF with FUS loss-of-function, IPF hLF, and IPF hLF with FUS gain-of-function. The resulting distributions were visualized using cumulative distribution plots.

### RNA polymerase II elongation speed analysis

RNA polymerase II elongation speed was estimated from RNA-seq data using the method described by Debès et al ^5^. The analysis was performed using the original scripts developed by the authors and made available through the Beyer laboratory repository.

High-quality RNA-seq reads were aligned to the human reference genome GRCh38/hg38, and read coverage was evaluated across intronic regions. Elongation dynamics were inferred from the decrease in nascent RNA-seq read coverage from the 5′ to the 3′ end of introns. For each retained intron, a linear regression was fitted to the normalized read-coverage profile, and RNA polymerase II elongation speed was estimated from the inverse relationship between the fitted coverage slope and transcriptional progression.

Elongation speed values were expressed as log_₁₀_(−1/slope). Introns with insufficient read coverage or unsuitable coverage profiles were excluded according to the filtering criteria implemented in the original analysis workflow. Elongation speeds were compared among Ctrl hLF, Ctrl hLF with FUS loss-of-function, IPF hLF, and IPF hLF with FUS gain-of-function. Average intronic coverage profiles and their corresponding fitted slopes were visualized for each experimental condition.

### Alternative splicing analysis

Alternative splicing events were analyzed from RNA-seq data using rMATS v4.1.2. High-quality reads were aligned to the human reference genome GRCh38/hg38 using STAR v2.7.9 with splice-junction detection enabled. The resulting BAM files were used as input for rMATS together with the GENCODE v39 gene annotation.

The analysis was performed in single-end mode using the parameters --gtf gencode.v39.annotation.gtf-t single --readLength 100 --nthread 8. Five major classes of alternative splicing events were evaluated: skipped exons, retained introns, alternative 5′ splice sites, alternative 3′ splice sites, and mutually exclusive exons.

For each event, rMATS calculated the percent spliced in value and assessed differential splicing between experimental conditions. Events with a P value < 0.05 and a minimum combined inclusion and skipping read count of 10 were retained for downstream analysis. Pairwise comparisons were performed between Ctrl hLF and Ctrl hLF with FUS loss-of-function, Ctrl hLF and IPF hLF, and IPF hLF and IPF hLF with FUS gain-of-function.

### Human Lung Cell Atlas analysis

Single-cell RNA-sequencing data were obtained from the Human Lung Cell Atlas through the CELLxGENE data portal. Cell-type annotations provided with the atlas were used to identify alveolar human lung fibroblasts and myofibroblasts. These populations were subset for downstream analyses of transcriptional trajectories, *FUS* expression, senescence-associated gene expression, and fibrotic signatures.

The cellular composition of the spatial niches reported by Mayr *et al.* 2024 ^9^ was used to guide the selection of corresponding populations from the single-cell RNA-sequencing atlas. Cell populations were selected according to their annotated identities and their correspondence with the dominant cellular components of the spatial niches.

Mayr *et al*. 2024 identified the fibrotic niche as an IPF-associated spatial compartment enriched in myofibroblasts, aberrant basaloid cells, basal cells, terminal bronchial secretory cells, and IL1B-positive macrophages. The airway macrophage niche was characterized by disease-associated macrophage populations, including CHI3L1-positive, CD9-high and C1Q-high macrophages, whereas the immune niche was enriched in B cells, T cells, dendritic cells, and PVLAP-positive bronchial endothelial cells ^9^.

Based on these spatial findings, atlas-annotated myofibroblasts were selected as the principal mesenchymal population representing the fibrotic niche. Alveolar human lung fibroblasts, annotated as alveolar fibroblasts in the single-cell atlas, were selected as a resident fibroblast population representative of the relatively non-fibrotic alveolar compartment. These two populations were independently subset for downstream analyses of pseudotime, *FUS* expression, fibrotic gene expression, and the SenMayo senescence signature.

To characterize transcriptional changes associated with pseudotime progression from Ctrl myofibroblasts toward IPF myofibroblasts, pseudotime trajectory analysis was performed using Monocle 3, with Ctrl myofibroblasts defined as the starting point of the trajectory ^49^. Cells with valid pseudotime values were retained, and gene expression was correlated with pseudotime using Spearman rank correlation. P values were adjusted for multiple testing using the Benjamini–Hochberg method, and genes with an FDR < 0.05 were considered significantly associated with pseudotime. Genes positively or negatively correlated with pseudotime were classified as increasing or decreasing, respectively. The expression of selected genes was standardized across cells using z-score transformation and visualized in a heatmap ordered according to pseudotime, with pseudotime and disease status (Control or IPF) included as annotations.

### Definition of FUS target genes by ChIP-seq and CLIP-seq

FUS target genes were defined by integrating FUS ChIP-seq and CLIP-seq datasets. First, genes associated with significant FUS enrichment in the ChIP-seq dataset and transcripts showing significant FUS binding in the CLIP-seq dataset were identified independently. Genes detected in both datasets were classified as FUS targets. The intersection yielded 7,150 genes, which were used for downstream analyses of R-loop enrichment and RNA polymerase II transcriptional dynamics.

### R-loop CUT&Tag

Genome-wide R-loop profiling was performed using the CUT&Tag-IT® R-loop Assay Kit (Active Motif, Cat. No. 53167), following the manufacturer’s instructions. Briefly, 2×10^6^ unfixed human lung fibroblasts from Ctrl hLF, Ctrl hLF with FUS loss-of-function and IPF hLF conditions were immobilized on concanavalin A-coated magnetic beads and permeabilized using the digitonin-containing buffers supplied with the kit. DNA-RNA hybrids were recognized using the kit-provided mouse monoclonal DNA-RNA hybrid antibody, clone S9.6. A rabbit anti-mouse secondary antibody was subsequently used to amplify recruitment of the adapter-loaded protein A-Tn5 transposome to the antibody-bound DNA-RNA hybrids. Following targeted tagmentation, DNA was released by proteinase K digestion and purified using the binding buffers and modified DNA-purification columns supplied with the kit. Sequencing libraries were amplified using the Q5 Polymerase Master Mix and indexed i5 and i7 primers included in the kit and were purified using SPRI beads. Library concentration, fragment-size distribution and integrity were evaluated before sequencing.

### R-loop CUT&Tag enrichment analysis in FUS targets

Sequencing adapters were trimmed, and high-quality paired-end reads were aligned to the human reference genome GRCh38/hg38 using Bowtie2 with the parameters --local --very-sensitive --no-mixed --no-discordant --phred33-I 10-X 700. SAM files were converted to BAM format using samtools view-Sb, and PCR duplicates were removed using Picard MarkDuplicates. Reads mapping to mitochondrial DNA or unassembled genomic sequences were excluded. Peaks were called using MACS2 v2.2.9.1 with the parameters callpeak-g hs-q 0.05 --broad --extsize 200 --keep-dup 1 --nomodel. Peaks overlapping ENCODE hg38 blacklisted regions were removed using bedtools intersect-v. RPKM-normalized genome-coverage tracks were generated from the filtered BAM files using bamCoverage from deepTools v3.5.6 and exported in BigWig format for downstream quantitative analyses and genome-browser visualization.

R-loop enrichment was subsequently evaluated at the 7,150 FUS target genes. TSS of the FUS targets was obtained from the hg38 gene annotation, and normalized S9.6 CUT&Tag signal was quantified ±2kb of the TSS. The Ctrl hLF R-loop profiles were subjected to unsupervised clustering, which identified four patterns according to the predominant position of R-loop enrichment relative to the TSS. Normalized R-loop CUT&Tag signal was visualized as average enrichment profiles and heatmaps and was quantitatively summarized for genes within each cluster. R-loop enrichment distributions were compared among experimental conditions using one-way analysis of variance followed by Tukey’s post hoc test for multiple comparisons. Genome-browser tracks were additionally generated in the Integrative Genomics Viewer to visualize the relationship between FUS enrichment, RNA polymerase II occupancy and R-loop formation at selected fibrotic and senescence-associated loci.

## Availability of data and materials

The data supporting this study are provided with this paper. Source data are provided with this paper as a Source Data file. The sequencing data generated in this study have been deposited in NCBI’s Gene Expression Omnibus database under accession numbers GSE349855, GSE349592. Proteomics raw data, processing and statistical analysis can be accessed at the following repositories: Mass spectrometry-based proteomics data: ProteomeXchange Consortium via the MassIVE partner repositories (http://proteomecentral.proteomexchange.org) with the following identifiers: ProteomeXchange: PXD083326, MassIVE ID: MSV000103037. These data sets remain private during manuscript peer review and will become publicly available upon manuscript acceptance for publication. Private access token for peer reviewers can be obtained from the journal editor. Furthermore, we retrieved and used publicly available datasets to aid analysis of our data. Extended Data Table 3 contains all data sets used in this study.

## Acknowledgements

We thank Roswitha Bender and Anne Robert for technical support; Virgine Marchand, Iouri Motorine and the EpiRNA-Seq Core Facility for support with NGS-based methods; Kerstin Richter, Alessandro Ianni, Sylvain Maenner and Bruno Charpentier for reagents; Sandrine Gulberti, Jean-Baptiste Vincourt, Sylvie Fournel-Gigleux, Mohamed Ouzzine, Catherine Bui, Lydia Barré and Nick Ramalanjaona for comments.

## Authors contributions

DGRA, HAAA, KR, GS, NC, NDI, WS, GK, VR, PB, SM, SG, JC and GB designed and performed the experiments; MNR, JG, TB, MW and GD were involved in study design; GB, and JC designed the study; DGRA, JC, HAAA, NC, NDI, WS, SG, JG, MW, GD and GB analyzed the data; GB, DGRA, HAAA, WS, JG, MW, and JC wrote the manuscript. All authors discussed the results and commented on the manuscript.

## Extended data figure/table legends

**Extended Data Fig. 1:**
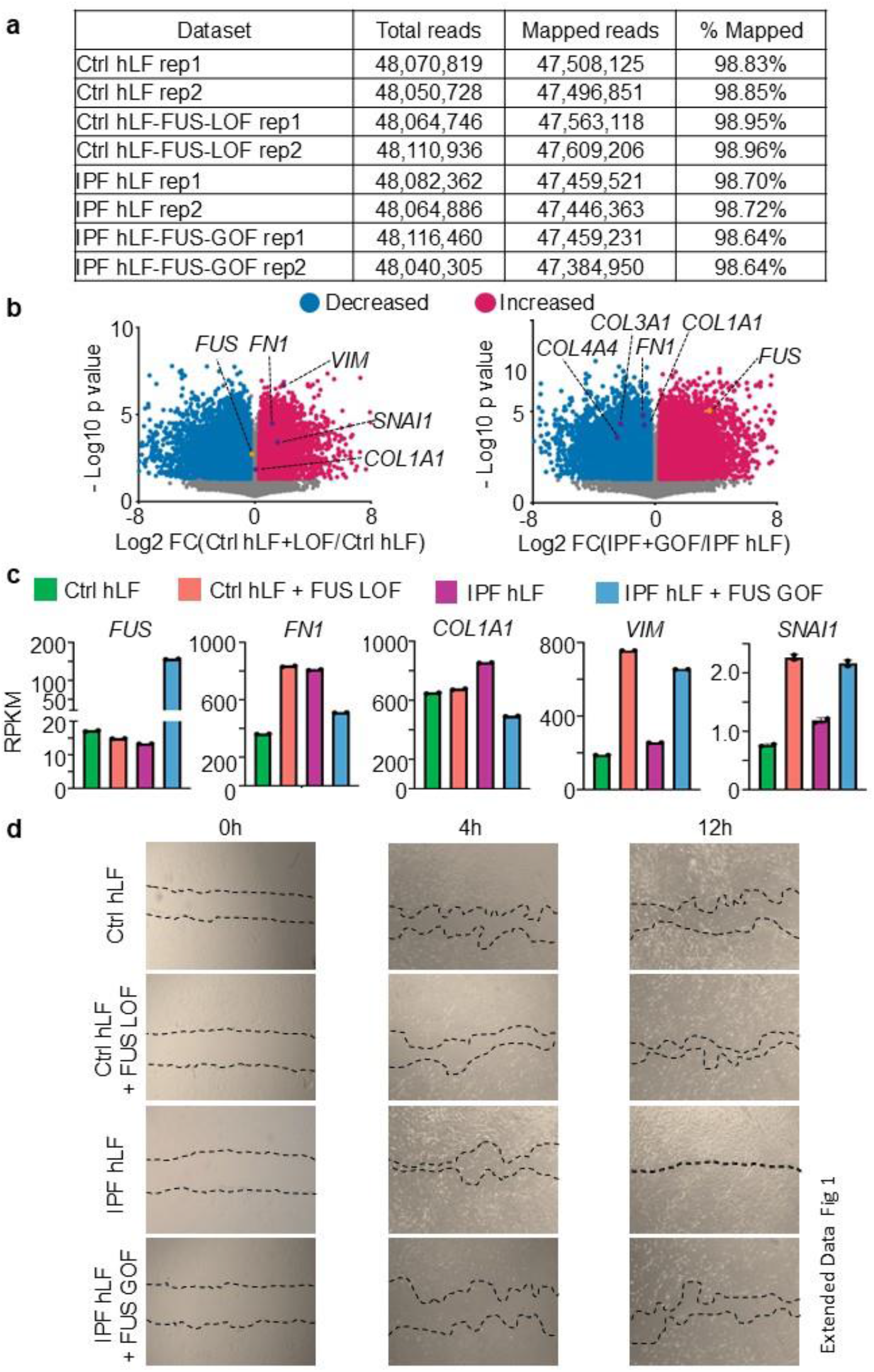
FUS modulates profibrotic gene expression and cell migration. **a**, RNA-seq read and mapping statistics for Ctrl hLF, Ctrl hLF + FUS LOF, IPF hLF and IPF hLF + FUS GOF samples. **b**, Volcano plots showing differentially expressed genes in Ctrl hLF + FUS LOF *versus* Ctrl hLF (left) and IPF hLF + FUS GOF *versus* IPF hLF (right). Genes with decreased and increased expression are shown in blue and magenta, respectively. Selected differentially expressed genes are labelled. **c**, Expression levels of FUS and the indicated fibrotic markers across the four experimental conditions, expressed as reads per kilobase per million mapped reads (RPKM). **d**, Representative phase-contrast images of scratch-wound assays performed in Ctrl hLF, Ctrl hLF + FUS LOF, IPF hLF and IPF hLF + FUS GOF at 0, 4 and 12 h. Dotted lines indicate the wound margins. See also Fig. 1. Source data are provided as a Source Data file.

**Extended Data Fig. 2:**
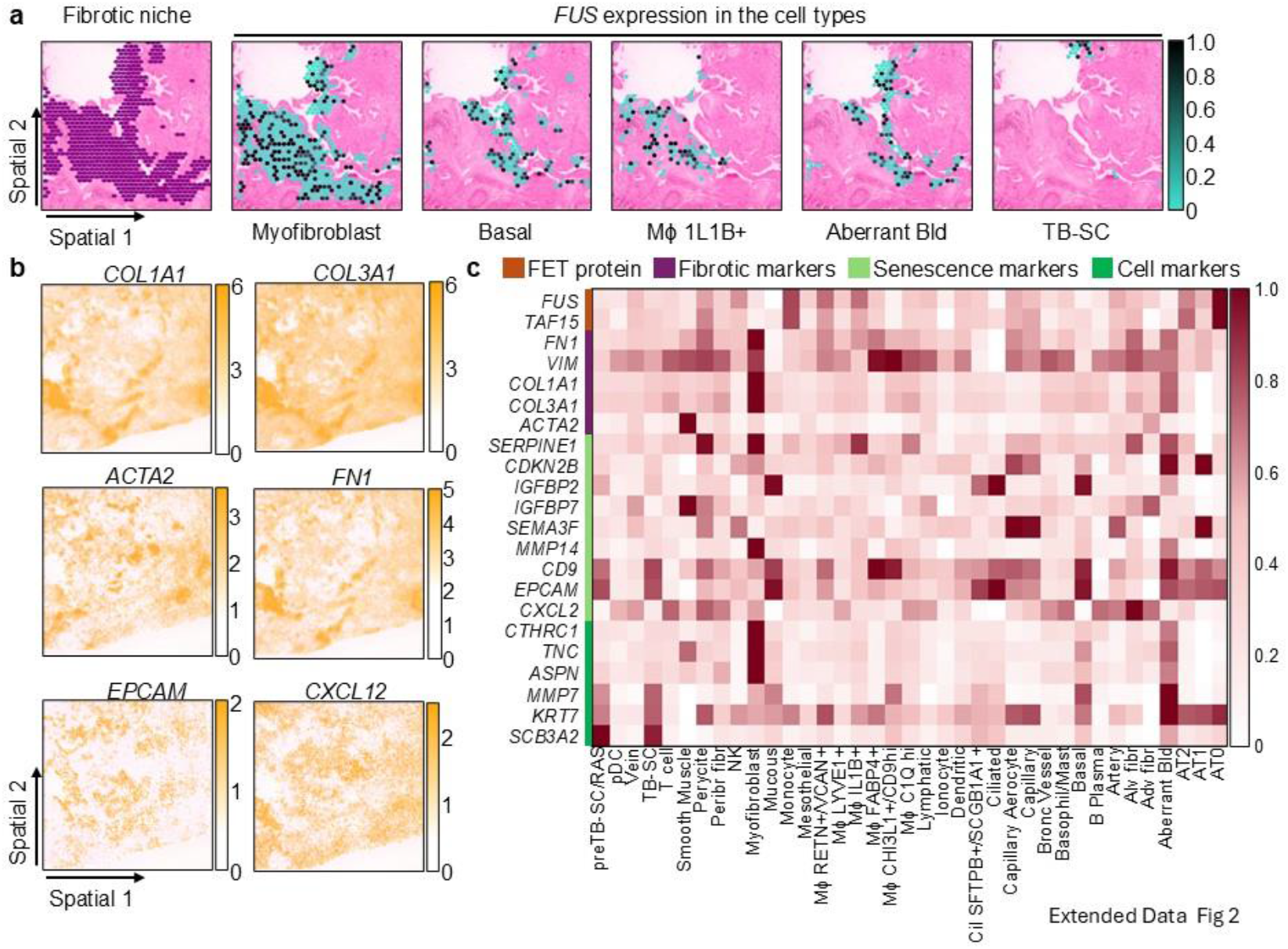
Spatial characterization of cell populations within the IPF fibrotic niche. **a**, Spatial transcriptomic maps showing the fibrotic niche and FUS expression in myofibroblasts, basal cells, IL1B+ macrophages, aberrant basaloid cells and terminal bronchial secretory cells (TB-SCs). **b**, Spatial expression maps of selected fibrotic and cell-type marker genes in the same IPF lung section. **c**, Heatmap showing the normalized expression of FUS, fibrotic markers, senescence markers and cell-type markers across annotated lung cell populations. See also Fig. 2. Source data are provided as a Source Data file.

**Extended Data Fig. 3:**
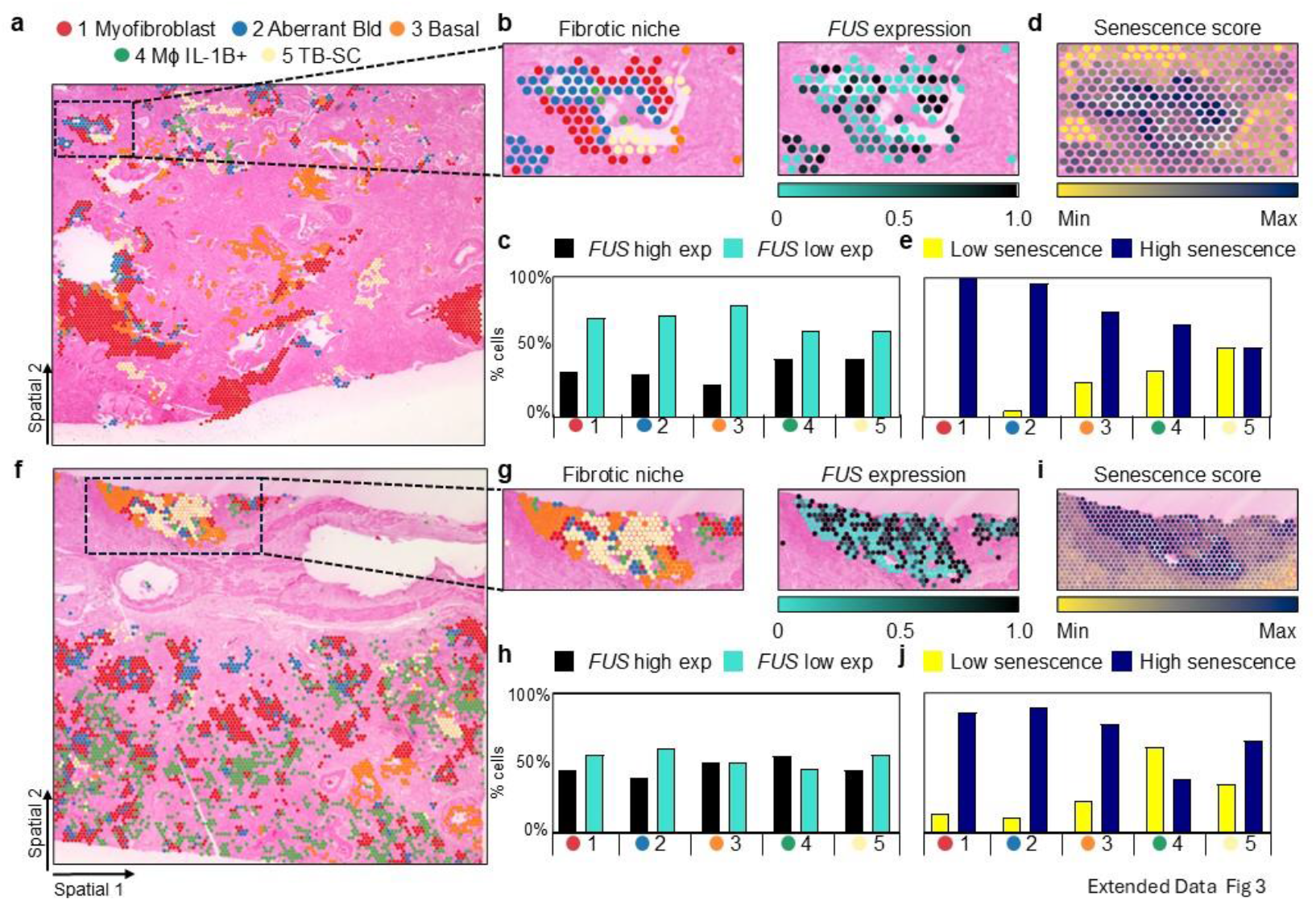
Low FUS expression coincides with high senescence scores in additional IPF lung sections. **a** and **f**, Spatial transcriptomic map showing the distribution of myofibroblasts, aberrant basaloid cells, basal cells, IL1B+ macrophages and terminal bronchial secretory cells (TB-SCs) in two different IPF lung sections. The box indicates the fibrotic niche magnified in b and d. **b** and **g**, Magnified view of the indicated fibrotic niche showing the selected cell populations and FUS expression. Dashed lines delineate the fibrotic niche. **c** and **h**, Percentages of cells with high or low FUS expression within each cell population. **d** and **i**, Spatial distribution of senescence scores in the indicated fibrotic niche. **e** and **j**, Percentages of cells with low or high senescence scores within each cell population. See also Fig. 2. Source data are provided as a Source Data file.

**Extended Data Fig. 4:**
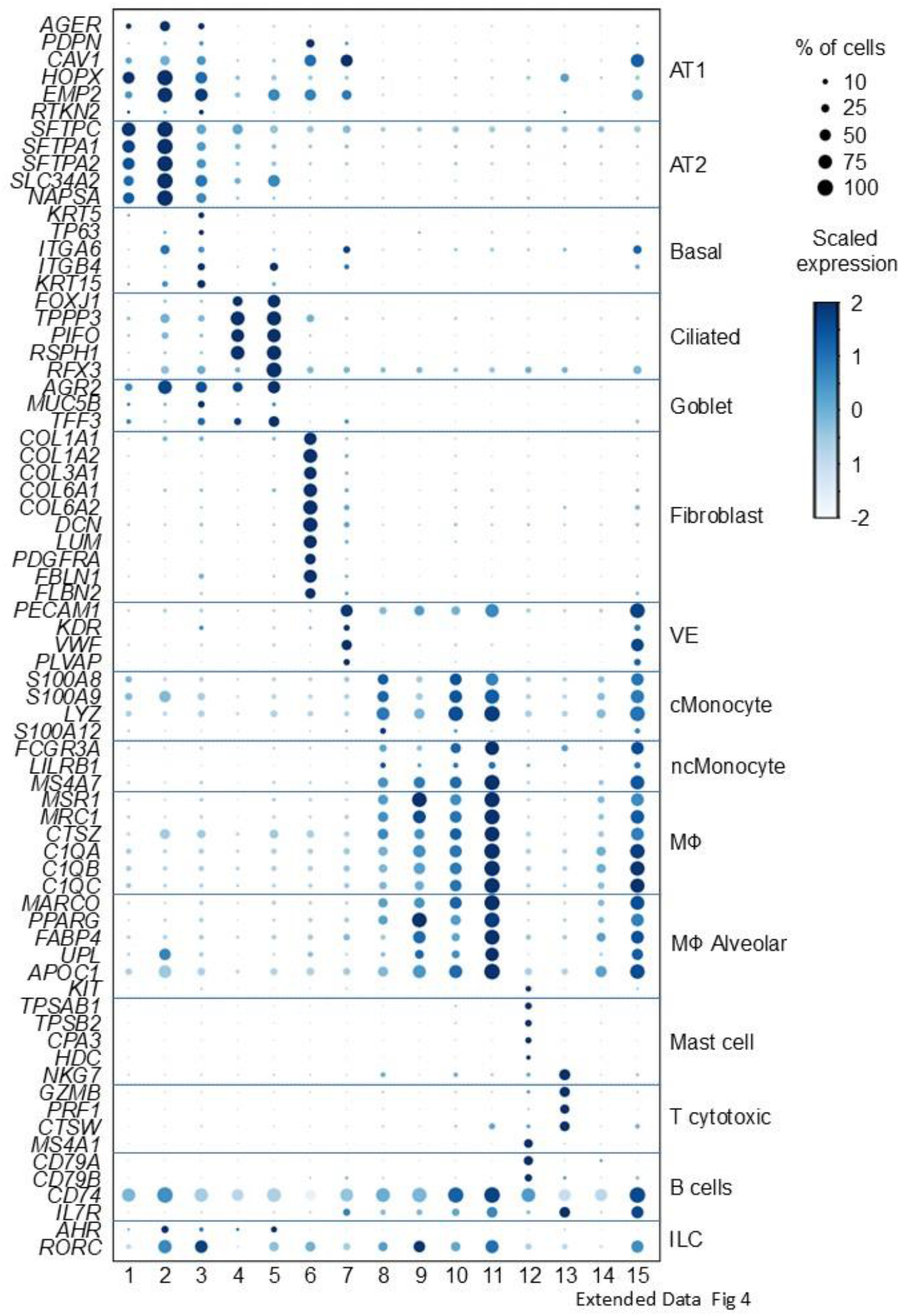
Dot plot showing the expression of cell-type marker genes across 15 annotated cell populations from Ctrl and IPF lungs. Dot size indicates the percentage of cells expressing each gene, and colour indicates scaled expression. VE, vascular endothelial cells; cMonocyte, classical monocytes; ncMonocyte, non-classical monocytes; Mφ, macrophages; Mφ Alveolar, alveolar macrophages; ILC, innate lymphoid cells. See also Fig. 3. Source data are provided as a Source Data file.

**Extended Data Fig. 5:**
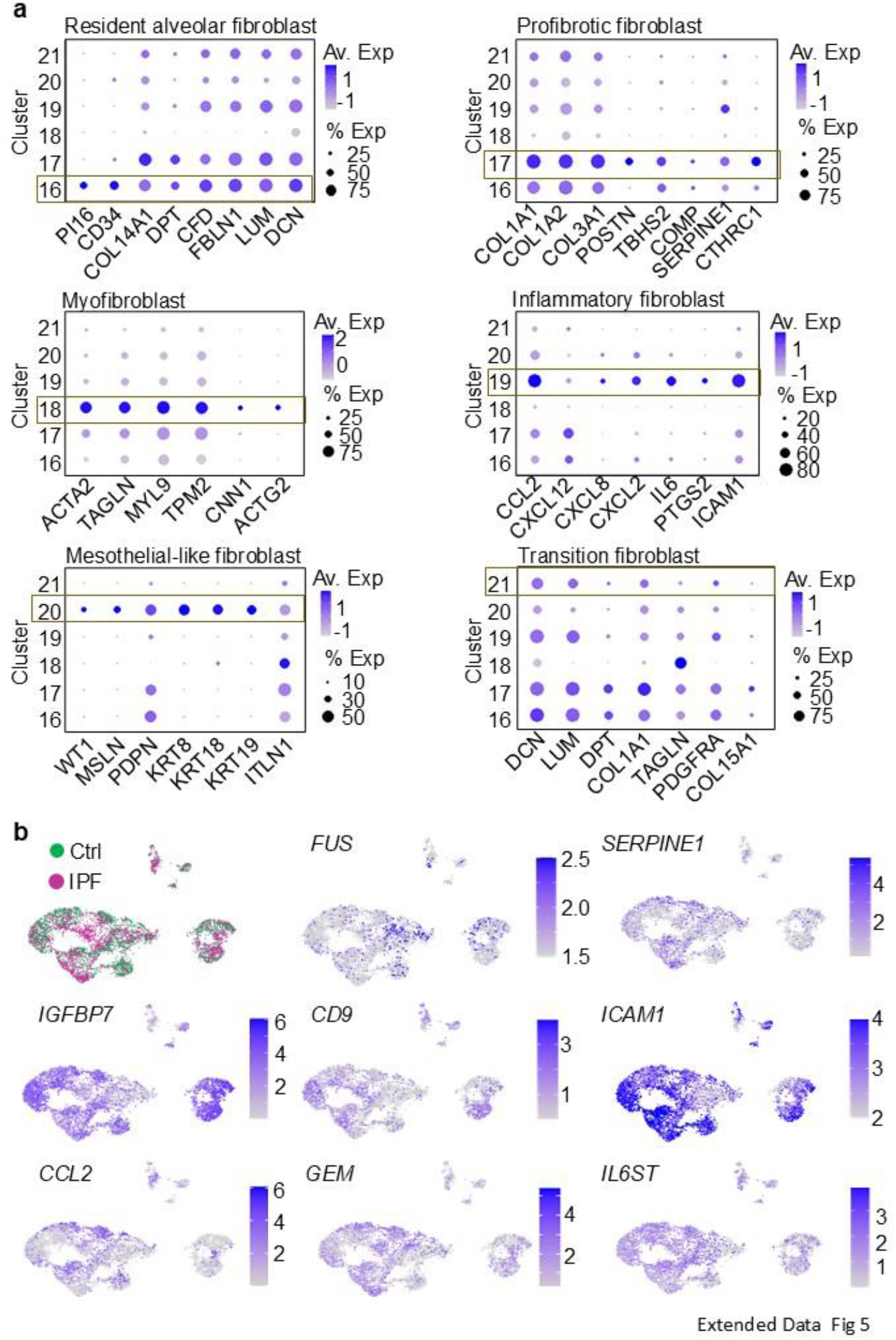
**Marker-gene profiles define six fibroblast subtypes in Ctrl and IPF lungs**. **a**, Dot plots showing the expression of marker genes used to annotate resident alveolar, profibrotic, myofibroblast, inflammatory, mesothelial-like and transition fibroblasts. Yellow boxes indicate the cluster assigned to each fibroblast subtype. Dot size indicates the percentage of cells expressing each gene, and colour indicates average expression. **b**, UMAP visualization of fibroblasts coloured by condition (Ctrl, green; IPF, magenta) and feature plots showing the normalized expression of FUS and selected fibroblast marker genes. See also Fig. 3. Source data are provided as a Source Data file.

**Extended Data Fig. 6:**
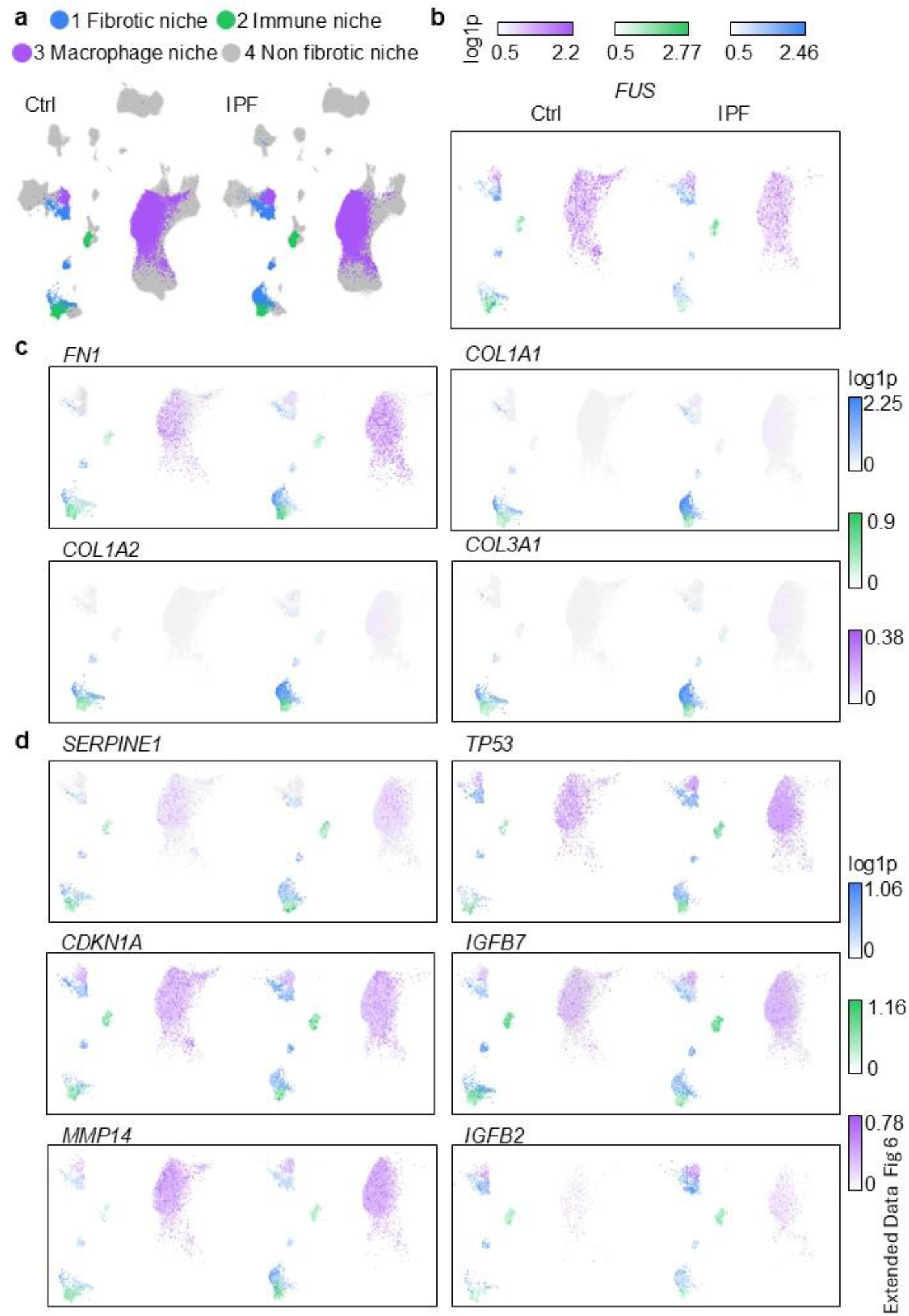
Cellular niches display distinct FUS, fibrotic and senescence-associated gene-expression specific patterns in Ctrl and IPF lungs. **a**, UMAP visualizations of Ctrl and IPF lung cells classified into fibrotic, immune, macrophage and non-fibrotic niches. **b**, UMAP feature plots showing FUS expression in Ctrl and IPF cells across the indicated niches. **c** and **d**, UMAP feature plots showing the expression of fibrosis-related genes (**c**) and senescence-associated genes (**d**) in Ctrl and IPF cells. Colour intensity indicates log1p-transformed expression. See also Fig. 4. Source data are provided as a Source Data file.

**Extended Data Fig. 7:**
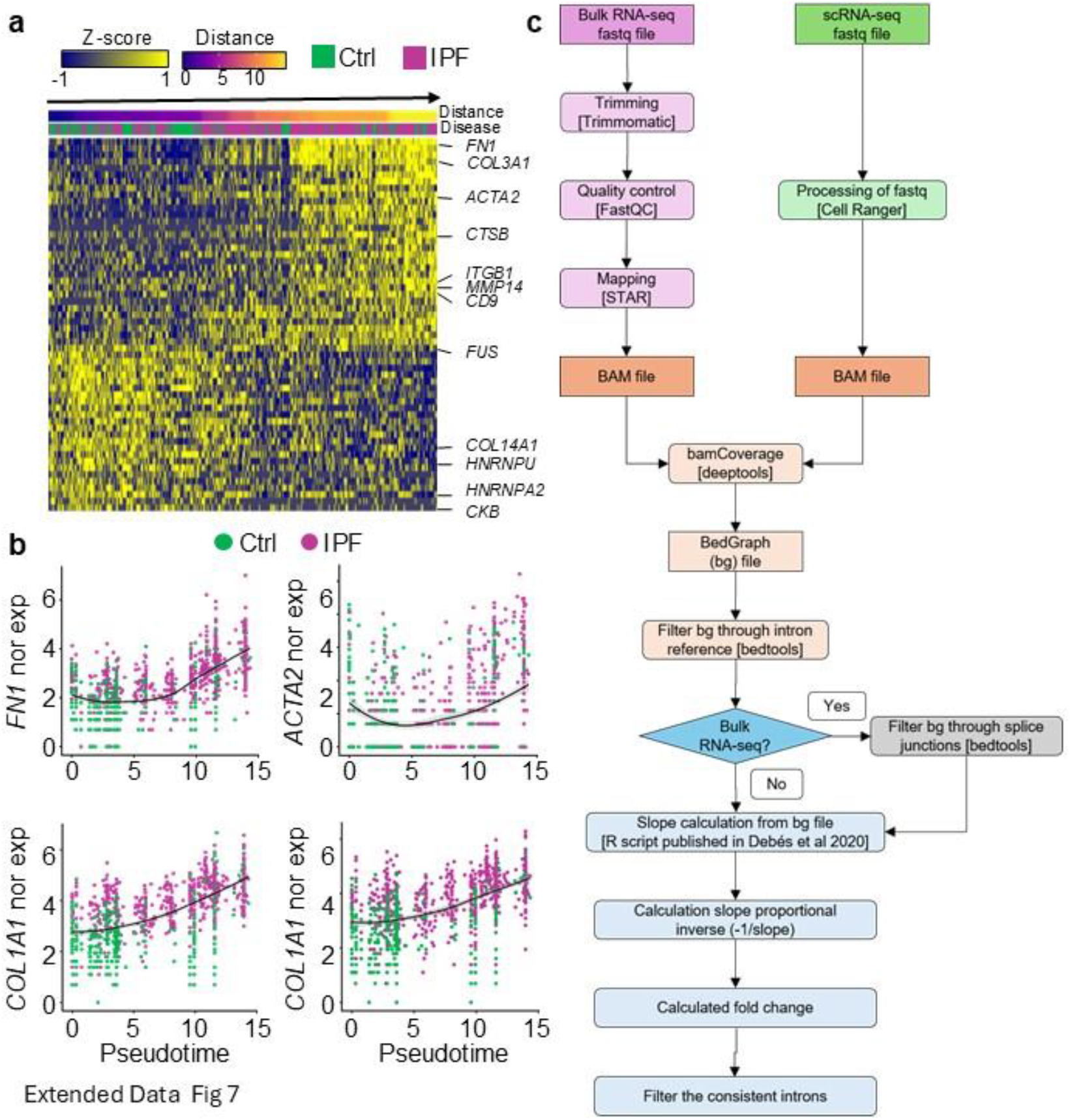
**Fibroblast pseudotime progression towards IPF is accompanied by increased expression of fibrotic genes**. **a**, Heatmap showing the z-scored expression of genes significantly correlated with diffusion pseudotime (DPT) in Ctrl and IPF myofibroblasts. Cells are ordered by pseudotime, with disease condition and DPT distance indicated by the top annotations. **b**, Smoothed expression of selected fibrosis-associated genes along pseudotime in Ctrl (green) and IPF (magenta) myofibroblasts. **c**, Schematic workflow for estimating cell type-specific Pol II elongation speed from intronic RNA-seq read coverage. See also Fig. 4. Source data are provided as a Source Data file.

**Extended Data Fig. 8:**
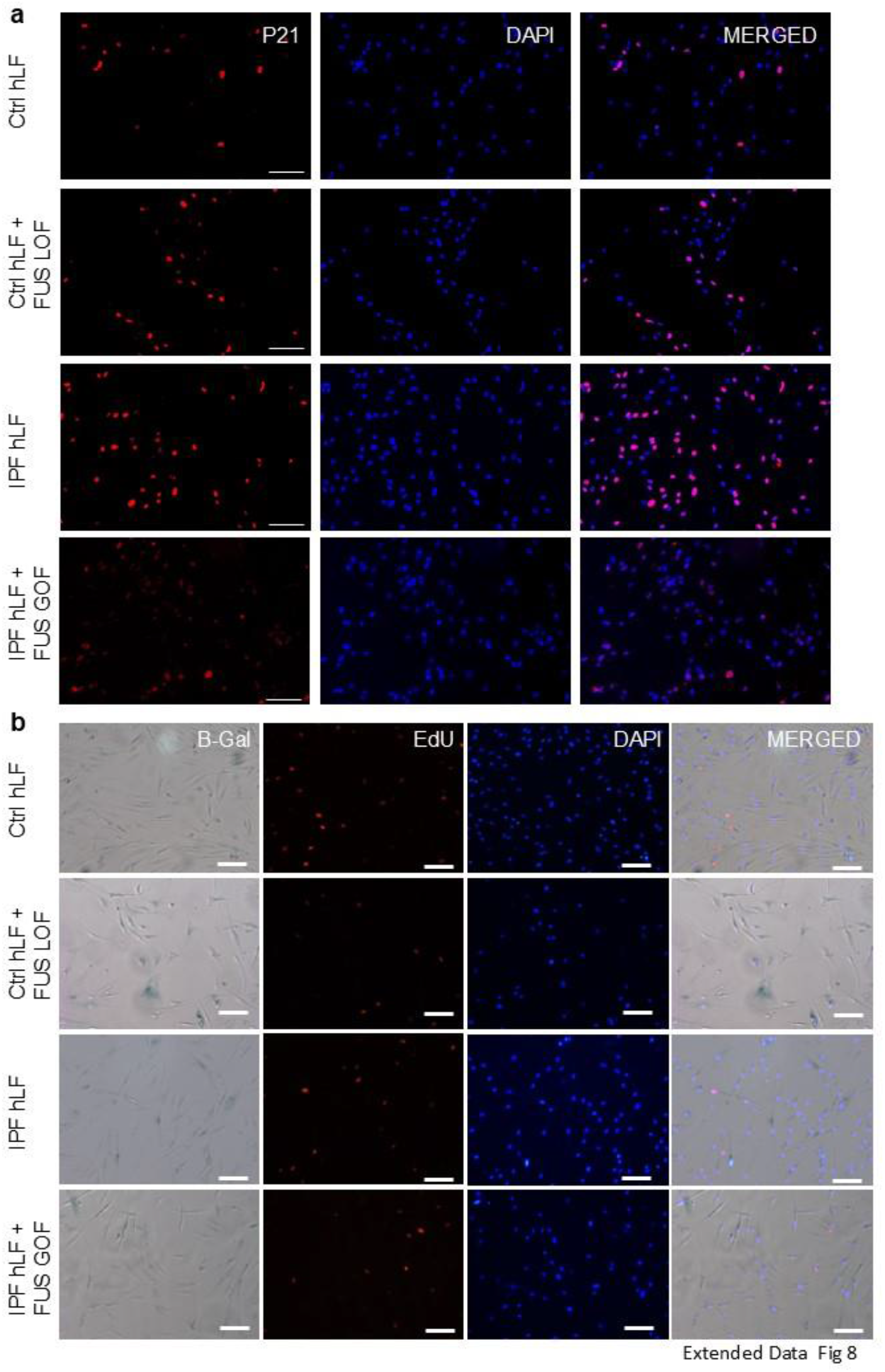
FUS loss promotes senescence-associated phenotypes in Ctrl hLF. **a**, Representative immunofluorescence images showing p21 (red) and DAPI-stained nuclei (blue) in Ctrl hLF, Ctrl hLF + FUS LOF, IPF hLF and IPF hLF + FUS GOF. **b**, Representative images showing senescence-associated β-galactosidase (SA-β-gal) staining, EdU incorporation (red), DAPI-stained nuclei (blue) and merged images in the indicated conditions. Scale bars are shown in the images. See also Fig. 5. Source data are provided as a Source Data file.

**Extended Data Fig. 9:**
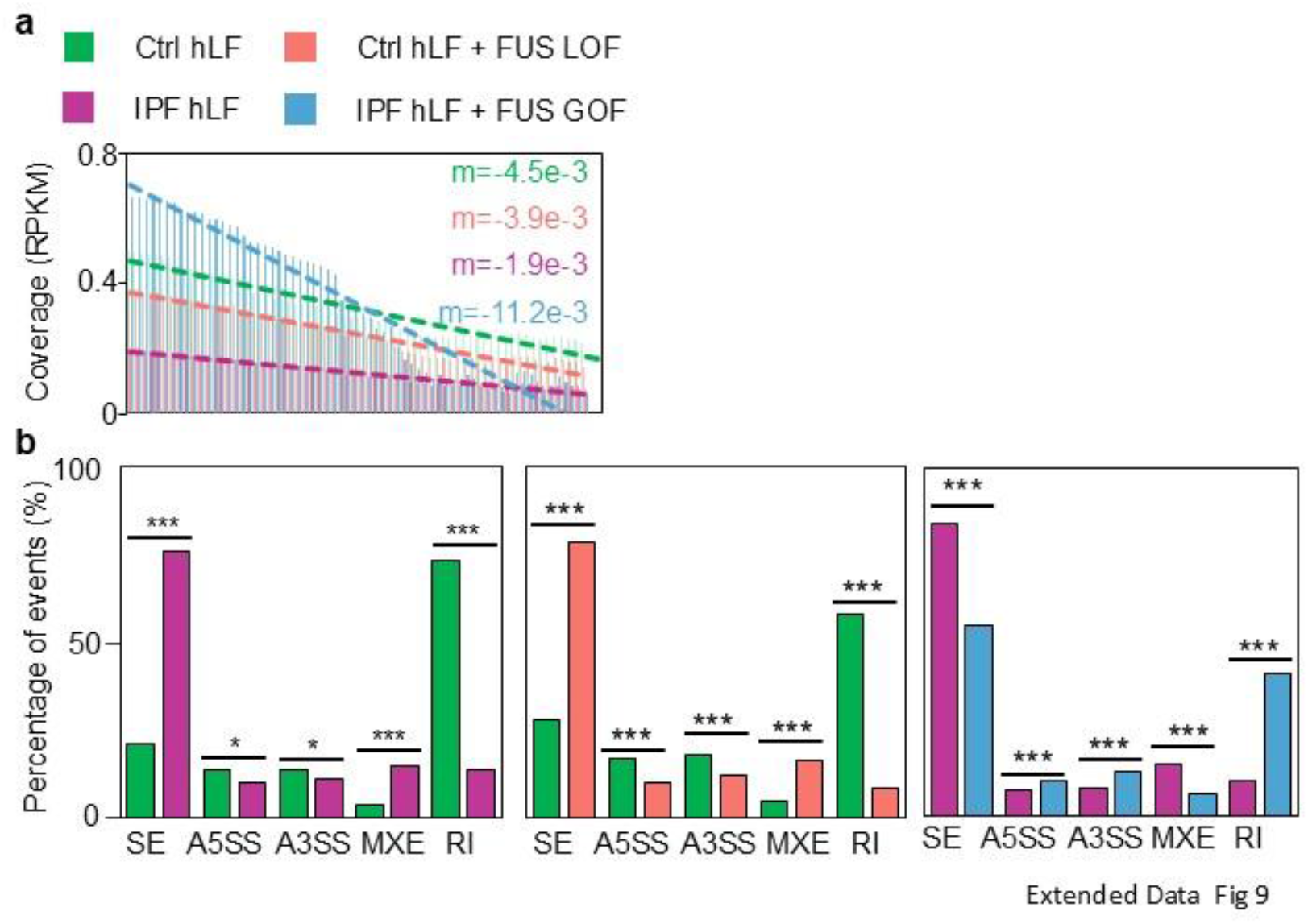
FUS loss increases Pol II elongation speed and alters alternative splicing patterns in hLF. **a**, Average intronic RNA-seq coverage profiles and fitted regression slopes (m) in Ctrl hLF, Ctrl hLF + FUS LOF, IPF hLF and IPF hLF + FUS GOF. The fitted slopes were used to estimate Pol II elongation speed. **b**, Percentages of the indicated alternative splicing event classes in IPF hLF versus Ctrl hLF, Ctrl hLF + FUS LOF versus Ctrl hLF, and IPF hLF + FUS GOF versus IPF hLF. SE, skipped exon; A5SS, alternative 5′ splice site; A3SS, alternative 3′ splice site; MXE, mutually exclusive exon; RI, retained intron. Bars represent mean. Statistical significance was assessed by two-tailed Student’s t-tests; *P < 0.05, **P < 0.01, ***P < 0.001. See also Fig. 5. Source data are provided as a Source Data file.

**Extended Data Fig. 10:**
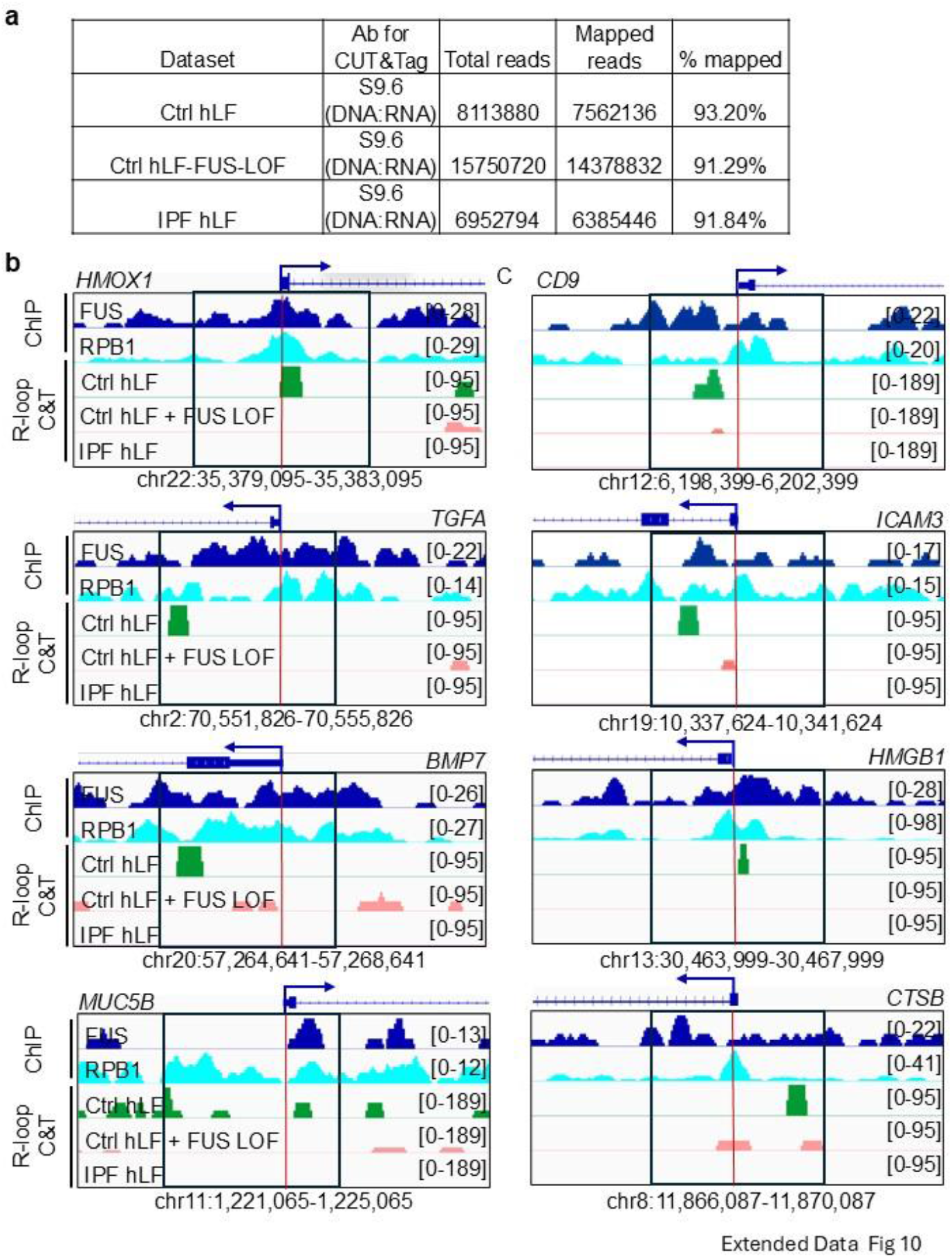
**Reduced FUS levels diminish R-loop enrichment at selected fibrosis-and senescence-associated loci**. **a**, Sequencing and mapping statistics for S9.6 R-loop CUT&Tag in Ctrl hLF, Ctrl hLF + FUS LOF and IPF hLF. **b**, Genome browser tracks showing FUS and RPB1 ChIP-seq signals together with R-loop CUT&Tag signals in Ctrl hLF, Ctrl hLF + FUS LOF and IPF hLF at the indicated loci. Red vertical lines indicate transcription start sites, and signal ranges are shown on the right. See also Fig. 6. Source data are provided as a Source Data file.

---

**Extended Data Table 1:**
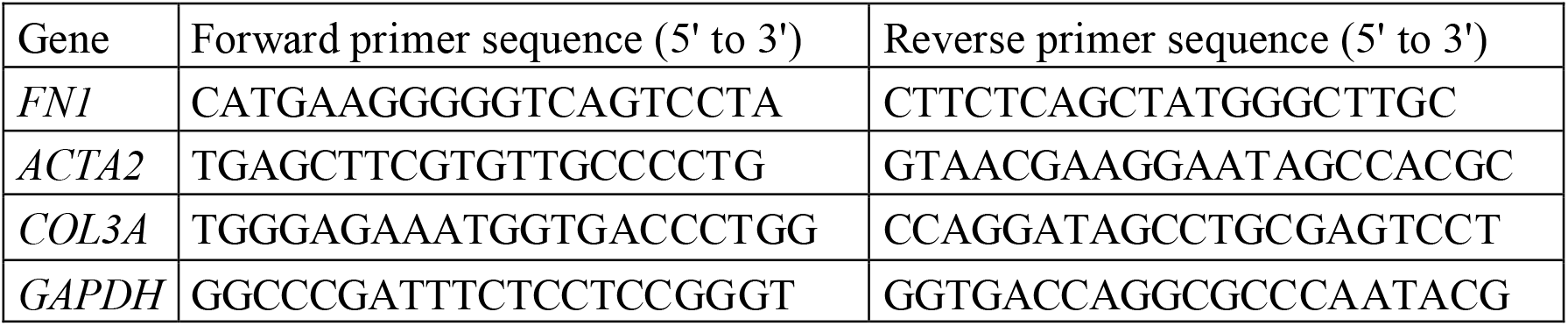
Primers used for ChIP-qPCR.

**Extended Data Table 2:** Antibodies used for WB, ChIP, CUT&TAG and IF.

| **Antibody** | **Company** | **Cat. No** | **WB** | **ChIP** | **CUT&Tag** | **IF** |
| --- | --- | --- | --- | --- | --- | --- |
| FUS | Santa Cruz | 47711 | YES<br>(1:1000) | YES | NO | YES<br>(1:500) |
| NMP1 | Santa Cruz | 56622 | NO | NO | NO | YES<br>(1:200) |
| RPB1 | Abcam | ab5408 | NO | YES | NO | NO |
| S9.6 | Kerafast | ENH001 | NO | NO | YES | NO |
| FN1 | BD | 610077 | YES<br>(1:1000) | NO | NO | NO |
| A-SMA | Biozol | GTX124505-50 | YES<br>(1:1000) | NO | NO | YES<br>(1:200) |
| COL1A1 | Rockland | 600-401-103 | YES<br>(1:1000) | NO | NO | NO |
| EXOSC10 | Abcam | 50558 | YES<br>(1:1000) | NO | NO | NO |
| GAPDH | Cell signalling | 97166 | YES<br>(1:1000) | NO | NO | NO |
| Goat anti-Rabbit IgG Alexa Fluor 488 | Thermofisher Scientific | A-11008 | NO | NO | NO | YES<br>(1:1000) |
| Goat anti-Mouse IgG Alexa Fluor 594 | Thermofisher Scientific | A-11005 | NO | NO | NO | YES<br>(1:1000) |
| rIgG | Santa Cruz | sc-2027 | YES<br>(1:1000) | YES | NO | NO |
| mIgG | Santa Cruz | sc-2025 | YES<br>(1:1000) | YES | NO | NO |
| gIgG | Santa Cruz | sc-2028 | YES<br>(1:1000) | NO | NO | NO |

**Extended Data Table 3.** Publicly available data sets used in this study.

| Dataset | NGS Method | Database | Dataset ID |
| --- | --- | --- | --- |
| Processed count object of Visium spatial transcriptomics data from Ctrl and IPF patients | Visium | Zenodo | https://zenodo.org/records/10012934 |
| Processed counts object of the integrated PF-ILD atlas from publicly available scRNA-seq datasets. | scRNA-seq | Zenodo | https://zenodo.org/records/10015169 |
| Ctrl_001C | scRNA-seq | GEO | GSM4058919 |
| Ctrl_208C | scRNA-seq | GEO | GSM4058910 |
| Ctrl_218C | scRNA-seq | GEO | GSM4058903 |
| Ctrl_226C | scRNA-seq | GEO | GSM4058904 |
| Ctrl_092C | scRNA-seq | GEO | GSM4058913 |
| Ctrl_001C | scRNA-seq | GEO | GSM4058919 |
| Ctrl_483C | scRNA-seq | GEO | GSM4058918 |
| Ctrl_484C | scRNA-seq | GEO | GSM4058924 |
| Ctrl_034C | scRNA-seq | GEO | GSM4058902 |
| Ctrl_084C | scRNA-seq | GEO | GSM4058927 |
| Ctrl_244C | scRNA-seq | GEO | GSM4058905 |
| Ctrl_465C | scRNA-seq | GEO | GSM4058907 |
| Ctrl_222C | scRNA-seq | GEO | GSM4058911 |
| Ctrl_160C | scRNA-seq | GEO | GSM4058912 |
| Ctrl_388C | scRNA-seq | GEO | GSM4058916 |
| Ctrl_137C | scRNA-seq | GEO | GSM4058926 |
| Ctrl_THD0001 | scRNA-seq | GEO | GSM4037313 |
| Ctrl_VUHD65 | scRNA-seq | GEO | GSM4037301 |
| Ctrl_VUHD66 | scRNA-seq | GEO | GSM4037302 |
| Ctrl_VUHD68 | scRNA-seq | GEO | GSM4037306 |
| Ctrl_VUHD70 | scRNA-seq | GEO | GSM4037316 |
| Ctrl_VUHD71 | scRNA-seq | GEO | GSM4037323 |
| Ctrl_SC156 | scRNA-seq | GEO | GSM6598824 |
| Ctrl_SC313 | scRNA-seq | GEO | GSM6598828 |
| Ctrl_SC315 | scRNA-seq | GEO | GSM6598829 |
| Ctrl_SC323 | scRNA-seq | GEO | GSM6598831 |
| Ctrl_SC333 | scRNA-seq | GEO | GSM6598833 |
| Ctrl_SC334 | scRNA-seq | GEO | GSM6598834 |
| Ctrl_SC45 | scRNA-seq | GEO | GSM6598817 |
| Ctrl_SC56 | scRNA-seq | GEO | GSM6598818 |
| IPF_40I | scRNA-seq | GEO | GSM4058968 |
| IPF_158I | scRNA-seq | GEO | GSM4058976 |
| IPF_177I | scRNA-seq | GEO | GSM4058977 |
| IPF_210I | scRNA-seq | GEO | GSM4058946 |
| IPF_214I | scRNA-seq | GEO | GSM4058956 |
| IPF_145I | scRNA-seq | GEO | GSM4058949 |
| IPF_221I | scRNA-seq | GEO | GSM4058957 |
| IPF_025I | scRNA-seq | GEO | GSM4058963 |
| IPF_47I | scRNA-seq | GEO | GSM4058969 |
| IPF_49I | scRNA-seq | GEO | GSM4058970 |
| IPF_TILD015 | scRNA-seq | GEO | GSM4037319 |
| IPF_VUILD61_! | scRNA-seq | GEO | GSM4037315 |
| IPF_VUILD65 | scRNA-seq | GEO | GSM4037322 |
| IPF_SC153 | scRNA-seq | GEO | GSM6598822 |
| IPF_SC154 | scRNA-seq | GEO | GSM6598823 |
| IPF_SC175 | scRNA-seq | GEO | GSM6598826 |
| IPF_SC307 | scRNA-seq | GEO | GSM6598827 |
| IPF_SC317 | scRNA-seq | GEO | GSM6598830 |
| IPF_SC93 | scRNA-seq | GEO | GSM6598820 |
| IPF_SC94 | scRNA-seq | GEO | GSM6598821 |
| FUS ChIP-seq in HEK293T | ChIP-seq | SRA | SRR642621 |
| Pol II ChIP-seq Ctrl in HEK293T | ChIP-seq | SRA | SRR646565 |

## References

1. Zhang, Q. et al. A tunable CTD grammar governs the spatial programming of the transcription cycle. Nat Commun 17, 6356 (2026).

2. Adelman, K. & Lis, J. T. Promoter-proximal pausing of RNA polymerase II: emerging roles in metazoans. Nat Rev Genet 13, 720–731 (2012).

3. Vos, S. M. et al. Structure of activated transcription complex Pol II-DSIF-PAF-SPT6. Nature 560, 607–612 (2018).

4. Schwartz, J. C. et al. FUS binds the CTD of RNA polymerase II and regulates its phosphorylation at Ser2. Genes Dev 26, 2690–2695 (2012).

5. Debès, C. et al. Ageing-associated changes in transcriptional elongation influence longevity. Nature 616, 814–821 (2023).

6. Noble, P. W., Barkauskas, C. E. & Jiang, D. Pulmonary fibrosis: patterns and perpetrators. J Clin Invest 122, 2756–2762 (2012).

7. Singh, I. et al. MiCEE is a ncRNA-protein complex that mediates epigenetic silencing and nucleolar organization. Nat Genet 50, 990–1001 (2018).

8. Rubio, K. et al. Inactivation of nuclear histone deacetylases by EP300 disrupts the MiCEE complex in idiopathic pulmonary fibrosis. Nat Commun 10, 2229 (2019).

9. Mayr, C. H. et al. Spatial transcriptomic characterization of pathologic niches in IPF. Sci Adv 10, eadl5473 (2024).

10. Habermann, A. C. et al. Single-cell RNA sequencing reveals profibrotic roles of distinct epithelial and mesenchymal lineages in pulmonary fibrosis. Sci Adv 6, eaba1972 (2020).

11. Adams, T. S. et al. Single-cell RNA-seq reveals ectopic and aberrant lung-resident cell populations in idiopathic pulmonary fibrosis. Sci Adv 6, eaba1983 (2020).

12. Kobayashi, Y. et al. Persistence of a regeneration-associated, transitional alveolar epithelial cell state in pulmonary fibrosis. Nat Cell Biol 22, 934–946 (2020).

13. Saul, D. et al. A new gene set identifies senescent cells and predicts senescence-associated pathways across tissues. Nat Commun 13, 4827 (2022).

14. DeTomaso, D. & Yosef, N. Hotspot identifies informative gene modules across modalities of single-cell genomics. Cell Syst 12, 446–456.e9 (2021).

15. Valenzi, E. et al. Single-nucleus chromatin accessibility identifies a critical role for TWIST1 in idiopathic pulmonary fibrosis myofibroblast activity. Eur Respir J 62, 2200474 (2023).

16. Haghverdi, L., Büttner, M., Wolf, F. A., Buettner, F. & Theis, F. J. Diffusion pseudotime robustly reconstructs lineage branching. Nat Methods 13, 845–848 (2016).

17. Ameur, A. et al. Total RNA sequencing reveals nascent transcription and widespread co-transcriptional splicing in the human brain. Nat Struct Mol Biol 18, 1435–1440 (2011).

18. Jonkers, I., Kwak, H. & Lis, J. T. Genome-wide dynamics of Pol II elongation and its interplay with promoter proximal pausing, chromatin, and exons. Elife 3, e02407 (2014).

19. Schuliga, M. et al. Mitochondrial dysfunction contributes to the senescent phenotype of IPF lung fibroblasts. J Cell Mol Med 22, 5847–5861 (2018).

20. Takebayashi, S.-I. et al. Retinoblastoma protein promotes oxidative phosphorylation through upregulation of glycolytic genes in oncogene-induced senescent cells. Aging Cell 14, 689–697 (2015).

21. Campisi, J. & d’Adda di Fagagna, F. Cellular senescence: when bad things happen to good cells. Nat Rev Mol Cell Biol 8, 729–740 (2007).

22. Dimri, G. P. et al. A biomarker that identifies senescent human cells in culture and in aging skin in vivo. Proc Natl Acad Sci U S A 92, 9363–9367 (1995).

23. Schwartz, J. C. et al. FUS is sequestered in nuclear aggregates in ALS patient fibroblasts. Mol Biol Cell 25, 2571–2578 (2014).

24. Rahl, P. B. et al. c-Myc regulates transcriptional pause release. Cell 141, 432–445 (2010).

25. Bentley, D. L. Coupling mRNA processing with transcription in time and space. Nat Rev Genet 15, 163–175 (2014).

26. Conaway, J. W. & Conaway, R. C. Transcription elongation and human disease. Annu Rev Biochem 68, 301–319 (1999).

27. Szklarczyk, D. et al. The STRING database in 2025: protein networks with directionality of regulation. Nucleic Acids Res 53, D730–D737 (2025).

28. Rubio, K., Dobersch, S. & Barreto, G. Functional interactions between scaffold proteins, noncoding RNAs, and genome loci induce liquid-liquid phase separation as organizing principle for 3-dimensional nuclear architecture: implications in cancer. FASEB J 33, 5814–5822 (2019).

29. Mosler, T. et al. R-loop proximity proteomics identifies a role of DDX41 in transcription-associated genomic instability. Nat Commun 12, 7314 (2021).

30. Cordero, J. et al. Nuclear microRNA 9 mediates G-quadruplex formation and 3D genome organization during TGF-β-induced transcription. Nat Commun 15, 10711 (2024).

31. Jutzi, D. et al. Aberrant interaction of FUS with the U1 snRNA provides a molecular mechanism of FUS induced amyotrophic lateral sclerosis. Nat Commun 11, 6341 (2020).

32. Yu, Y. & Reed, R. FUS functions in coupling transcription to splicing by mediating an interaction between RNAP II and U1 snRNP. Proc Natl Acad Sci U S A 112, 8608–8613 (2015).

33. Yang, L., Gal, J., Chen, J. & Zhu, H. Self-assembled FUS binds active chromatin and regulates gene transcription. Proc Natl Acad Sci U S A 111, 17809–17814 (2014).

34. Katariya, B. B. et al. Targeting fused in sarcoma (FUS): a novel antisense strategy for treating idiopathic pulmonary fibrosis. Signal Transduct Target Ther 11, 70 (2026).

35. Wang, I. X. et al. Human proteins that interact with RNA/DNA hybrids. Genome Res 28, 1405–1414 (2018).

36. Aguilera, P. & Aguilera, A. R-loop homeostasis in genome dynamics, gene expression and development. Curr Opin Genet Dev 92, 102325 (2025).

37. La Manno, G. et al. RNA velocity of single cells. Nature 560, 494–498 (2018).

38. Bergen, V., Soldatov, R. A., Kharchenko, P. V. & Theis, F. J. RNA velocity-current challenges and future perspectives. Mol Syst Biol 17, e10282 (2021).

39. Chamberlin, J. T., Lee, Y., Marth, G. T. & Quinlan, A. R. Differences in molecular sampling and data processing explain variation among single-cell and single-nucleus RNA-seq experiments. Genome Res 34, 179– 188 (2024).

40. He, D., Mount, S. M. & Patro, R. scCensus: Off-target scRNA-seq reads reveal meaningful biology. Preprint at 10.1101/2024.01.29.577807 (2024).

41. Butler, A., Hoffman, P., Smibert, P., Papalexi, E. & Satija, R. Integrating single-cell transcriptomic data across different conditions, technologies, and species. Nat Biotechnol 36, 411–420 (2018).

42. Wolf, F. A., Angerer, P. & Theis, F. J. SCANPY: large-scale single-cell gene expression data analysis. Genome Biol 19, 15 (2018).

43. Palla, G. et al. Squidpy: a scalable framework for spatial omics analysis. Nat Methods 19, 171–178 (2022).

44. Virshup, I. et al. The scverse project provides a computational ecosystem for single-cell omics data analysis. Nat Biotechnol 41, 604–606 (2023).

45. Kleshchevnikov, V. et al. Cell2location maps fine-grained cell types in spatial transcriptomics. Nat Biotechnol 40, 661–671 (2022).

46. Hughes, C. S. et al. Single-pot, solid-phase-enhanced sample preparation for proteomics experiments. Nat Protoc 14, 68–85 (2019).

47. Demichev, V., Messner, C. B., Vernardis, S. I., Lilley, K. S. & Ralser, M. DIA-NN: neural networks and interference correction enable deep proteome coverage in high throughput. Nat Methods 17, 41–44 (2020).

48. Ritchie, M. E. et al. limma powers differential expression analyses for RNA-sequencing and microarray studies. Nucleic Acids Res 43, e47 (2015).

49. Qiu, X. et al. Reversed graph embedding resolves complex single-cell trajectories. Nat Methods 14, 979–982 (2017).

